# Quantitative Proteomics and Phosphoproteomics Supports a Role for Mut9-Like Kinases in Multiple Metabolic and Signaling Pathways in *Arabidopsis*

**DOI:** 10.1101/2020.02.14.950030

**Authors:** Margaret E. Wilson, Shin-Cheng Tzeng, Megan M. Augustin, Matthew Meyer, Xiaoyue Jiang, Jae H. Choi, John C. Rogers, Bradley S. Evans, Toni M. Kutchan, Dmitri A. Nusinow

## Abstract

Protein phosphorylation is one of the most prevalent post-translational modifications found in eukaryotic systems. It serves as a key molecular mechanism that regulates protein function in response to environmental stimuli. The Mut9-Like Kinases (MLKs) are a plant-specific family of Ser/Thr kinases linked to light, circadian, and abiotic stress signaling. Here we use quantitative phosphoproteomics in conjunction with global proteomic analysis to explore the role of the MLKs in daily protein dynamics. Proteins involved in light, circadian, and hormone signaling, as well as several chromatin-modifying enzymes and DNA damage response factors, were found to have altered phosphorylation profiles in the absence of MLK family kinases. In addition to altered phosphorylation levels, *mlk* mutant seedlings have an increase in glucosinolate metabolism enzymes. Subsequently, we show that a functional consequence of the changes to the proteome and phosphoproteome in *mlk* mutant plants is elevated glucosinolate accumulation, and increased sensitivity to DNA damaging agents. Combined with previous reports, this work supports the involvement of MLKs in a diverse set of stress responses and developmental processes, suggesting that the MLKs serve as key regulators linking environmental inputs to developmental outputs.

## Introduction

Protein phosphorylation is a dynamic post-translational modification that is key in the regulation of protein function and turnover, making it an integral part of complex signaling networks. Rapid and reversible post-translational regulation is advantageous to plants, as they are often required to adapt to changing environments quickly. Moreover, protein phosphorylation is at the core of various biological processes, including stress response, light signaling, circadian regulation, and hormone perception and transduction. In Arabidopsis, nearly 4% of protein-encoding genes are kinases (Wang et al., 2014), which is a testament to the importance of phosphorylation-based protein regulation (Mergner et al., 2020). Despite the upswing of large-scale phosphoproteomic studies in plant species (Silva-Sanchez et al., 2015; Mergner et al., 2020), a recent study suggests that the identification of Arabidopsis phosphoproteins and phosphosites is far from comprehensive (Vlastaridis et al., 2017).

The four-member family of Ser/Thr protein kinases known as the <u>M</u>UT9-<u>l</u>ike <u>k</u>inase/<u>P</u>hotoregulatory <u>P</u>rotein <u>K</u>inases/<u>A</u>rabidopsis <u>EL</u>1-like (MLK/PPK/AEL) kinases, herein referred to as the MLKs, are involved in the phosphoregulation of several key signaling proteins (Ni et al., 2017; Liu et al., 2017; Su et al., 2017; Chen et al., 2018). The MLKs are a plant and green algae-specific family of kinases related to casein kinase I (CKI). The MLKs show significant divergence from CKI, with similarities restricted to their catalytic domains (Casas-Mollano et al., 2008). MLK family kinases are capable of phosphorylating histones H3 and H2A in *Arabidopsis* (Wang et al., 2015a; Kang et al., 2020) as well as the green algae *Chlamydomonas* (Casas-Mollano et al., 2008). The *Chlamydomonas* MLK orthologue, MUT9, is also required for transgene silencing and response to DNA damaging agents (Casas-Mollano et al., 2008; Jeong Br et al., 2002). In addition to phosphorylating histones, MLK family kinases phosphorylate proteins involved in multiple signaling pathways. Early studies of a rice MLK orthologue, EARLY FLOWERING1 (EL1), have linked this kinase family to hormone signaling and the regulation of flowering time (Dai and Xue, 2010), a role which is at least in part conserved in *Arabidopsis* (Zheng et al., 2017; Chen et al., 2018; Huang et al., 2016; Kang et al., 2020; Sun et al., 2020). MLKs also interact with core components of the morning (Zheng et al., 2017; Su et al., 2017) and evening (Huang et al., 2016) loops of the *Arabidopsis* circadian clock. The association of the MLKs with the evening complex components, EARLY FLOWERING 3 AND 4 (ELF3 and ELF4), is dependent on the presence of the red light receptor phytochrome B (Huang et al., 2016). Additionally, the MLKs phosphorylate the blue light receptor CRYPTOCHROME2 and the red light-regulated transcription factor PHYTOCHROME INTERACTING FACTOR 3 (Ni et al., 2017; Liu et al., 2017). In sum, these studies suggest that the MLKs provide a link between light and circadian signaling, which in turn regulates plant growth and development.

In this study, we used quantitative phosphoproteomic techniques to expand our understanding of the various signaling pathways and cellular protein networks regulated by the MLK family of kinases. We combined isobaric labeling with high pH reversed-phase prefractionation and TiO_2_ based phosphopeptide enrichment to achieve an in-depth phosphoproteomic analysis of wild-type and *mlk* mutant seedlings at two different time points, one at the end of the day (ZT12) and the other several hours into the night (ZT14). We identified over 20,000 phosphosites mapping to nearly 5,000 protein groups. Notably, MLK mutant seedlings have altered abundance of glucosinolate metabolism enzymes, and differential phosphorylation of proteins involved in a diverse set of biological processes, including RNA processing, chromatin organization, and stress responses. The confluence of stress and chromatin factors suggested that MLKs may also regulate DNA-damage responses in *A. thaliana*, which was tested by assessing the sensitivity of *mlk* mutants to DNA-damaging agents.

## Experimental Procedures

### Plant Material

The *mlk1* (SALK_002211; AT5G18190), *mlk2* (SALK_064333; AT3G03940), and *mlk3* (SALK_017102; AT2G25760) mutant lines were obtained from the ABRC (Ohio State University). The *mlk4* (GABI_756G08; AT3G13670) mutant line was obtained from the Nottingham Arabidopsis Stock Centre. All are in the Colombia (Col-*0*) background and were isolated as previously described (Huang et al., 2016).

### Tissue Collection for Mass Spectrometry

Arabidopsis wild type (Col-*0*) and mutant seedlings were grown on sterilized qualitative filter paper (Whatman) overlaid on ½ x MS (Murashige and Skoog) plates containing 1% sucrose and 0.8% agar at 22°C. Seedlings were entrained under 12 h white light (100-110 μmol/m^2^/s)/12 h dark cycle. Tissue was collected on the 10^th^ day of growth immediately before lights off [Zeitgeber 12, (ZT12)] or after 2h of dark (ZT14).

### Protein Isolation and Digestion

The seedlings were transferred into a liquid N2 chilled 35ml ball mill and disrupted in a reciprocal mixer mill [30 Hz, 45 seconds, repeated three times (Retsch USA)] under liquid nitrogen. Ground tissue was gently resuspended in 1 mL (approximately 1 packed tissue volume) of SII buffer (100 mM sodium phosphate, pH 8.0, 150 mM NaCl, 5 mM EDTA, 5 mM EGTA, 0.1% Triton X-100, 1 mM PMSF, 1x protease inhibitor cocktail [Roche], 1x Phosphatase Inhibitors II & III [Sigma], and 50 μM Mg-132 [Peptides International]) and sonicated twice at 40% power, 1 second on/off cycles for 20 s total on ice (Fisher Scientific model FB505, with microtip probe). Extracts were clarified by centrifugation twice at 4°C for 10 min at ≥20,000xg. Protein concentrations were determined by BCA protein assay (Thermo-Fisher Scientific, Rockford, IL). Protein samples were reduced with 10 mM TCEP and alkylated with 25 mM iodoacetamide before trypsin digestion in 1/40 enzyme/protein ratio at 37°C overnight.

### Phosphopeptide Enrichment

Phosphopeptide enrichment was performed using the High-Select™ TiO_2_ Phosphopeptide Enrichment kit (Thermo Scientific PN32993) following the vendor’s protocol. Briefly, dried peptides were reconstituted in 150 μL of binding/equilibration buffer provided and applied to the TiO_2_ spin that was previously equilibrated with binding buffer/equilibration. After reapplying sample once, the tip was sequentially washed twice with 20 μL of binding buffer and wash buffer, and once with 20 μL of LC-MS grade water. Bound peptides were eluted by two applications of 50 μL of elution buffer (also provided). Eluates containing the enriched phosphopeptides were dried down and subsequently resuspended with 50 μl 0.1% formic acid for peptide concentration measurement using the Pierce Quantitative Colorimetric Assay kit (Thermo Scientific PN23275).

### Tandem Mass Tag (TMT) Labeling

100 μg of each digested sample was added to 100 μL of 100 mM HEPES pH 8.5 buffer. A reference pooled sample composed of equal amounts of material from all samples was also generated to link TMT experiments. Isobaric labeling of the samples was performed using 10-plex tandem mass tag (TMT) reagents (Thermo Fisher Scientific, Rockford, IL). All individual and pooled samples were labeled according to the TMT 10-plex reagent kit instructions. Briefly, TMT reagents were brought to room temperature and dissolved in anhydrous acetonitrile. Peptides were labeled by the addition of each label to its respective digested sample. Labeling reactions were incubated for 1 h at room temperature. Reactions were terminated with the addition of hydroxylamine.

### High pH Reverse Phase Fractionation

High pH reverse phase fractionation was performed using the Pierce High pH Reversed-Phase Peptide Fractionation Kit (Thermo Scientific PN84868) according to the manufacturer’s instructions. Briefly, peptide samples were dissolved in 300 μL of 0.1% TFA solution in LC-MS grade water and subsequently loaded onto reversed-phase fractionation spin columns also equilibrated with 0.1% TFA. Samples were then washed with 300 μL of 5% ACN/ 0.1% TEA to remove unreacted TMT reagent. Peptides were eluted into 8 peptide fractions with an ACN step gradient (i.e. 10%, 12.5%, 15%, 17.5%, 20%, 22.5%, 25%, and 50%). Samples were acidified and dried down prior to LC-MS.

### LC-MS/MS Analysis

Two microliters (one microgram) of each sample was injected onto a 0.075 x 500 mm EASY-Spray Pepmap C18 column equipped with a 0.100 x 5 mm EASY-Spray Pepmap C18 trap column (Thermo-Fisher Scientific, San Jose, CA) attached to an EASY-nLC 1000 (Thermo-Fisher Scientific, San Jose, CA). The peptides were separated using water (A) and acetonitrile (B) containing 0.1% formic acid as solvents at a flow rate of 300 nL per minute with a three-hour gradient. Data were acquired in positive ion data-dependent mode on an Orbitrap Fusion Lumos mass spectrometer (Thermo-Fisher Scientific, San Jose, CA) with a resolution of 120,000 (at *m/z* 200) and a scan range from *m/z* 380-1500. Precursor isolation was performed using the quadrupole prior to either CID activation in the ion trap and detection in the Orbitrap at a resolution of 30,000 or HCD activation with detection in the Orbitrap at a resolution of 60,000.

### Data Analysis

All MS/MS data were analyzed using Proteome Discoverer 2.1 (Thermo-Fisher Scientific, San Jose, CA). The search algorithm used in the study was Byonic v2.11 as part of the Proteome Discoverer software platform. Precursor ion mass tolerance was set to 10 ppm, and fragment ion tolerance was 20 ppm; up to 2 missed cleavages were allowed. Carbamidomethylation (+57.021 Da) on cysteine and TMT tag (+229.163 Da) on peptide N-termini as well as lysine residues were set as static modifications. Dynamic modifications included acetylation (+42.011 Da) on protein N-termini, oxidation (+15.995 Da) on methionine and phosphorylation (+79.966 Da) on serine, threonine, and tyrosine. Data were searched against the TAIR10 database (20101214, 35,386 entries) with FDR set to 1%.

For quantitation, reporter ion intensity integration tolerance was set to 20 ppm. Reporter ion abundances were corrected for isotopic impurities based on manufacturer’s specifications. For each peptide, a minimal average reporter S/N threshold of 2 and a co-isolation threshold of 100% are required. The S/N values for all peptides were summed within each TMT channel, and each channel was scaled according to the reference channel. Both unique and razor peptides were used for quantification.

Peptides of altered abundance were identified from the Byonic output list generated from HCD MS2 analysis using Microsoft Excel. Abundance ratios for mutant/wild type pairwise comparisons were calculated from the average peptide abundance of mutant and wild-type biological replicates. Only peptides identified in at least two biological replicates were considered for further analysis. Statistical significance was determined by Student’s *t*-test (*P*-value ≤ 0.05).

### Bioinformatic Analysis

The Motif-X algorithm (Chou and Schwartz, 2011) was used to extract significantly enriched phosphorylation motifs from *mlk1/2/3* and *mlk1/3/4* phosphopeptide data sets. Only phosphopeptides with high confidence phosphorylation sites were used in the analysis. The peptides were aligned and extended to a width of 15 amino acids using the online utility PEPTIDEXTENDER ver.0.2.2 alpha (schwartzlab.uconn.edu/pepextend/). The aligned peptides were used to extract motifs. The probability threshold was set to p-value ≤ 10^−5^; the occurrence threshold was set to 10. The default IPI Arabidopsis Proteome data set was used as the background data set.

Enrichment analysis of Gene ontology (GO) categories was performed with g:Profiler (Reimand et al., 2016). AGI accession numbers for Arabidopsis were uploaded to the g:Profiler web server (http://biit.cs.ut.ee/gprofiler/), and GO enrichment was determined using default settings (significance level 0.05). Enriched terms were summarized, and redundancy removed using the online tool REVIGO (Supek et al., 2011). Semantic similarity threshold (dispensability) was set to 0.5 (default) for all global proteome analysis and cellular component category of the phosphoproteome analysis. Dispensability was increased to 0.7 for all other analyses.

### Glucosinolate Extraction and Analysis by HPLC and LC-MS/MS

Arabidopsis seeds (Col-0, *mlk 1/2/3*, and *mlk 1/3/4*) were sown on ½ x MS (Murashige and Skoog) plates containing 1% sucrose and 0.8% agar and grown under 12 h white light (100-110 μmol/m^2^/s)/12 h dark cycle at 22°C for 10 days before harvesting at ZT12. Glucosinolates were extracted from approximately 350 mg of whole seedlings and desulfonated (in quadruplicate) as previously described (Crocoll *et al*. 2016) using sinigrin as an internal standard. Desulfo-GLS extracts were analyzed by HPLC (Waters) equipped with a photodiode array detector and separated using a Gemini C-18 column (150 × 2.00 mm, 5 μm; Phenomenex) with a flow rate of 0.5 mL per minute and the following solvents and binary gradient: solvent A-water and solvent B-acetonitrile; where solvent B was held at 1.5% for 1 min, then 1-6 min 1.5-5% B, 6-8 min 5-7% B, 8-18 min 7-21% B, 18-23 min 21-29% B, 23-30 min 29-43% B, 30-33 min 43-100% B, 33-37 100% B, 37-38 min 100-1.5% B, and held at 1.5% B for an additional 7 minutes. GLS peaks were identified using previously published UV spectra in addition to reported relative retention times and quantitated using peak areas of desulfo-GLS and internal standard along with published response factors (Brown *et al*. 2003, Grosser and Dam 2017). GLS identities were confirmed by LC-MS/MS (SCIEX 6500 QTRAP, Framingham, MA) using enhanced product ion (EPI; ion trap MS/MS) scans to verify the presence of previously published fragment ions (Crocoll et al. 2016) from each glucosinolate ion. Mass spectrometric data were collected in positive ion mode using the same gradient/solvents/column as for HPLC-UV analysis with the following source conditions: curtain gas, 20; ion-spray voltage, 5500 V; temperature, 500° C; gas 1, 40; gas 2, 45; declustering potential, 80 V; entrance potential, 10 V; collision energy 20 eV.

### MMS Treatment

Arabidopsis wild type (Col-*0*) and mutant seed was surface-sterilized and sown on ½ x MS (Murashige and Skoog) plates containing 1% sucrose and 0.8% agar with or without methyl methanesulfonate (MMS, Sigma). After stratification for two days at 4°C, seedlings were grown under 12 h white light (100-110 μmol/m^2^/s)/12 h dark cycle at 22°C. For growth sensitivity assays, seedlings were germinated on control media, and after five days of growth were transferred to MMS treatment media. Fresh weight was measured after 15 days of growth in the presence of MMS. For post-germination developmental assessment, the seed was germinated on ½ x MS plates containing 1% sucrose, 0.8% agar, and 150 ppm MMS. Seedlings were imaged and scored for arrest at 12 days after germination.

### UV-C Tolerance Assay

Whole-plant sensitivity to UV-C (254 nm) was evaluated as described in Castells et al. (2010) with the following modifications. 8-day old seedlings were irradiated with 2000 or 4000 J m^−2^ of UV-C twice during a 48 hr time period using a Stratalinker^®^ UV Crosslinker 1800. Following each treatment, plants were returned to growth conditions of 12 h white light (100-110 μmol/m^2^/s)/12 h dark cycle at 22°C. Seedlings were imaged after five days of recovery, and their phenotypes were measured.

## Results

### Proteomic Analysis Reveals Changes in Stress Response Pathways in *mlk* Mutant Seedlings

We measured the regulatory effects of the MLKs on proteome dynamics by tandem mass tag (TMT) labeling combined with high pH reversed-phase fractionation and tandem mass spectrometry (**Fig. 1**). Wild-type and *mlk* mutant *Arabidopsis* seedlings were entrained under 12 h light and 12 h dark conditions. We compared *mlk1/2/3* and *mlk1/3/4* mutant seedlings, as we were unable to isolate viable *mlk1/2/3/4* mutant seed (Huang et al., 2016; Liu et al., 2017). This mutant combination will allow us to assess potential redundancy within the MLK family and facilitate the identification of *mlk2* or *mlk4* specific changes. As the MLKs are associated with light-signaling pathways, we collected tissue either immediately before lights off [Zeitgeber 12, (ZT12)] or after 2h of dark (ZT14). We identified nearly 50,000 peptides combined, mapping to over 7,500 protein groups at both ZT12 and ZT14. Pairwise comparisons between *mlk* mutants and wild type identified peptides showing altered abundance in the *mlk* mutant backgrounds at both the ZT12 and ZT14 time-points (**Supplemental Dataset S1**). Peptides were classified as altered in abundance if both the log2 FC was at least ±1 (2 fold-change) and the p-value < 0.05. Only 13 unique proteins met our altered abundance criteria in the *mlk1/2/3* mutant when compared to wild type at both ZT12 and ZT14 (**Fig. 2 and Supplemental Table S1**). In the *mlk1/3/4* mutant background, more than 225 peptides mapping to over 110 unique proteins met our altered abundance threshold at ZT12 and ZT14 when compared to wild type (**Fig. 2 and Supplemental Table S1**). These results suggest that the *mlk1/3/4* mutant combination has a greater impact on the global proteome than the *mlk1/2/3* mutant combination at the light-to-dark transition.

**Figure 1.**
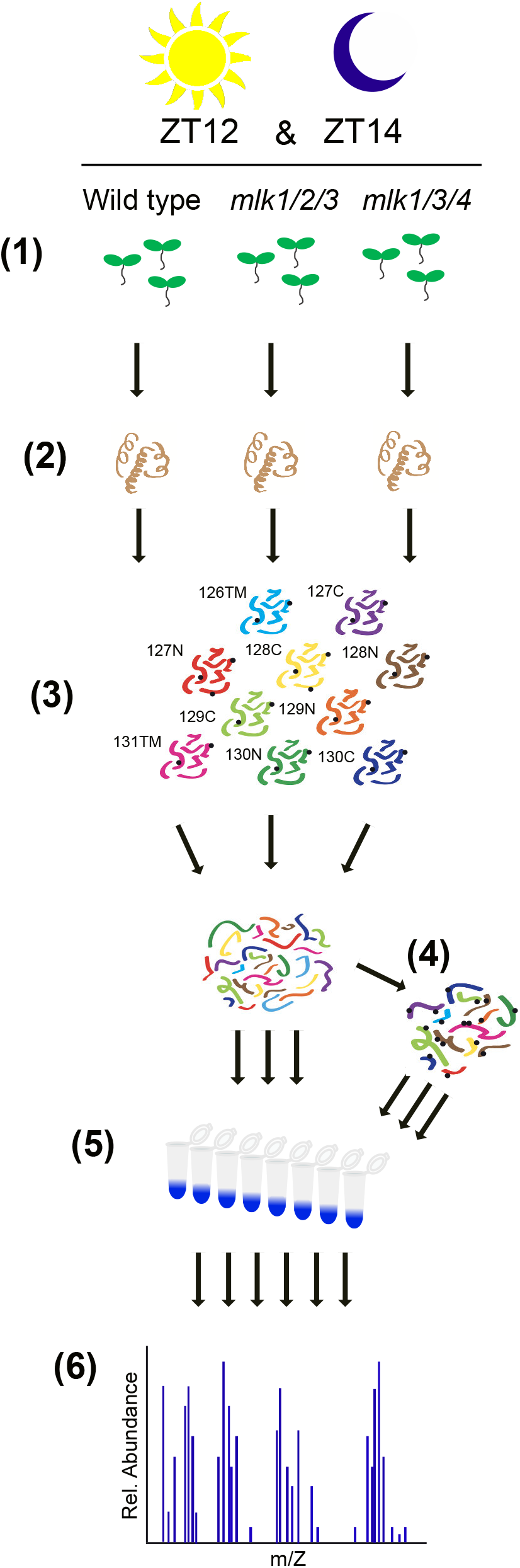
Schematic Representation of the Quantitative Proteomics Workflow. Tissue samples were collected at ZT12 and ZT14 from wild-type and mutant seedlings entrained with a 12L:12D light/dark cycle (1). Total protein was extracted and digested (2). Following TMT10plex isobaric labeling (3), samples were subjected to high pH reversed phase prefractionation (5). Phosphopeptides were enriched using a TiO_2_-based method (4). Both phosphopeptide-enriched and global samples were analyzed by LC-MS/MS (6).

**Figure 2.**
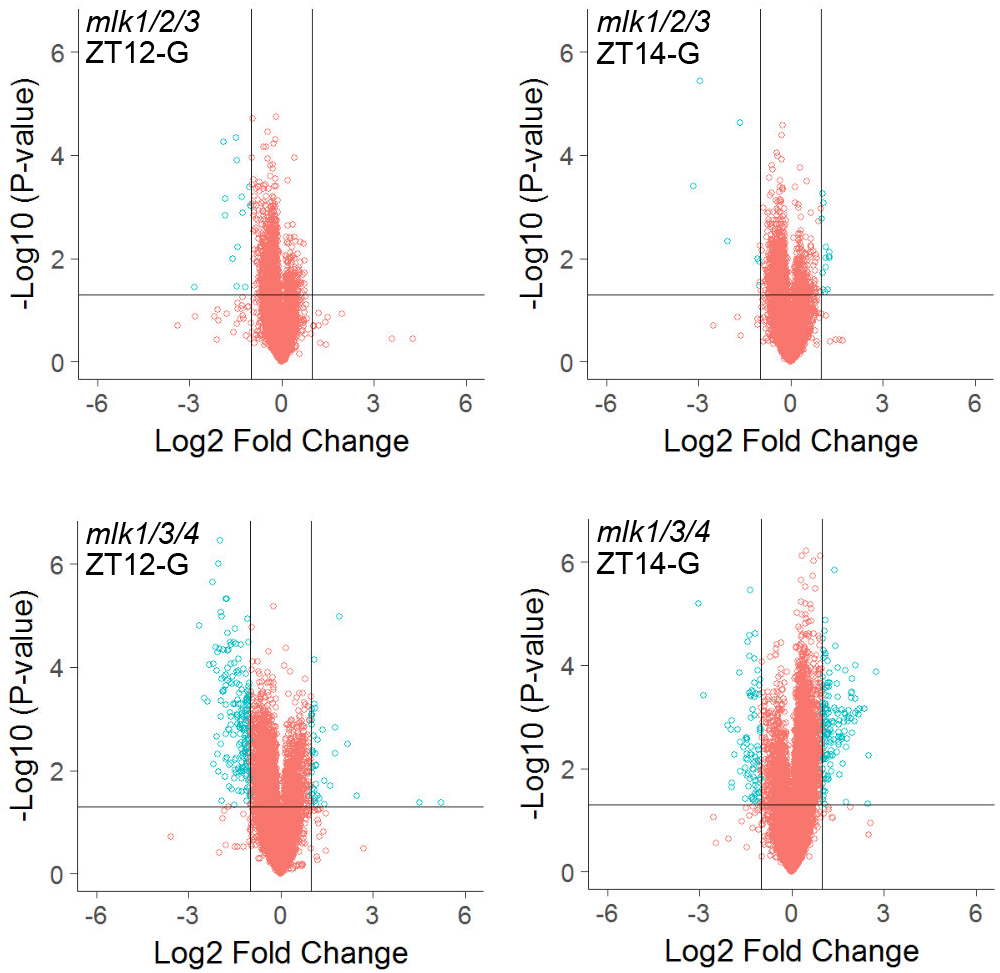
Global Proteomic Analysis of *mlk* Mutant Seedlings. Volcano plots of peptides identified in mutant and wild-type seedlings at ZT12 and ZT14. The x-axis specifies the log2 fold-change (FC) of mutant/wild type and the y-axis specifies the negative logarithm to the base 10 of the t-test p-values. Open circles represent individual peptides, with blue circles specifying those considered statistically significant. Black vertical and horizontal lines reflect the filtering criteria (log2 FC = ±1 and p-value = 0.05) for significance.

*In silico* classification using Gene Ontology (GO) analysis (https://biit.cs.ut.ee/gprofiler/) revealed that proteins exhibiting altered abundance were associated with biotic and abiotic stresses (**Fig. 3**). To simplify the enriched GO term lists and focus on the most relevant terms, we performed additional analysis using REVIGO (default settings, dispensability threshold = 0.7) to remove functionally redundant terms (Supek et al., 2011). We found increased abundance of proteins involved in glucosinolate biosynthesis (GO:0019761) and related processes (GO:1901659, GO:0016143, and GO:0044272) in *mlk1/3/4* mutant seedlings at ZT12 and in both *mlk1/2/3* and *mlk1/3/4* mutant seedlings at ZT14 (**Fig. 3**). Nearly 75% of the peptides with an increased abundance of 3-fold or greater in the *mlk1/3/4* mutant seedlings at ZT14 mapped to proteins directly involved in glucosinolate biosynthesis (**Supplemental Dataset S1**). These proteins include enzymes responsible for the early reactions leading to methionine-derived glucosinolates (branched-chain aminotransferase 4 (BCAT4) and methylthioalkymalate synthase 1 (MAM1)) as well as, desulfo-glucosinolate sulfotransferase 17 and 18 (SOT17 and SOT18), which are involved in the final step of glucosinolate core structure biosynthesis (Sønderby et al., 2010). Other proteins involved in glucosinolate biosynthesis that showed increased abundance in *mlk1/3/4* mutants compared to wild type include isopropymalate dehydrogenase 1 (IMD1), iospropylmaate isomerase 2 (IPMI2), 2-isopropylmalate synthase 2 (IMS2),flavin-monooxygenase glucosinolate S-oxygenase 1 (FMO GS-OX1) and the cytochrome P450 proteins CYP83A1 and CYP79F1 (**Supplemental Dataset S1**). Proteins involved in glucosinolate catabolism, such as glucoside glucohydrolase 2 (TGG2), nitrile specifier protein 1 (NSP1), and beta glucosidase 34 and 35 (BGLU34 and BGLU35) were decreased in abundance in *mlk1/3/4* mutant seedlings at ZT12 compared to wild type.

**Figure 3.**
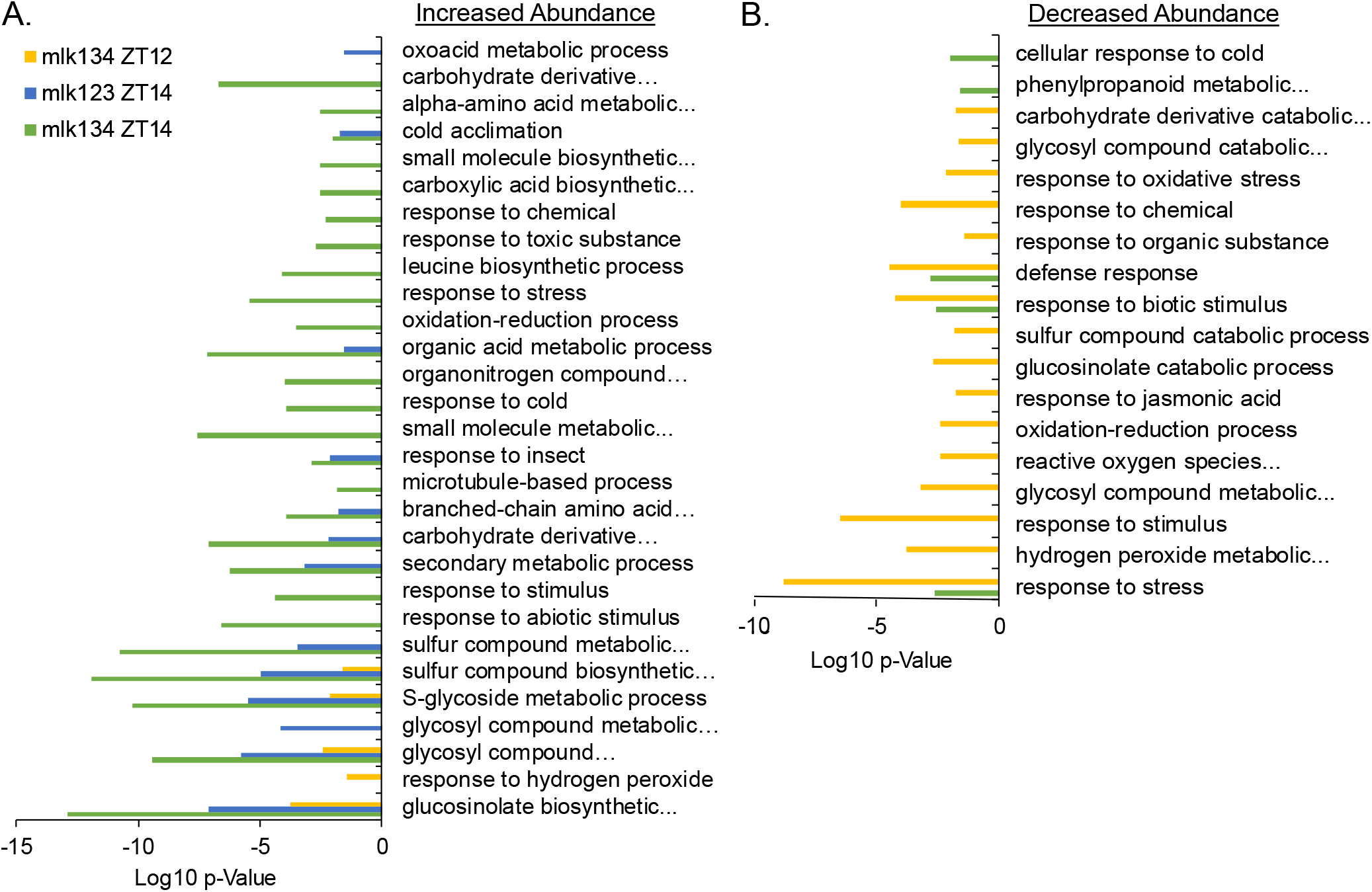
Gene Ontology (GO) Enrichment Analysis. GO enrichment analysis of proteins identified as having increased (A) or decreased (B) abundance in *mlk* mutant seedlings at indicated ZTs when compared to wild type. Cluster representative GO terms identified with REVIGO (semantic similarity threshold < 0.7) in the category of Biological Process are shown.

To begin testing the hypothesis that MLKs are involved in the regulation of glucosinolate metabolism, we quantified glucosinolate levels at the end of day (ZT12) when GLS levels peak (Huseby et al., 2013). Total GLSs were extracted from whole seedlings and analyzed by HPLC. Peaks corresponding to individual GLSs were identified by comparison with published UV absorbance spectra and expected retention times, and the identities were further validated using LC-MS/MS. These analyses revealed an increase in aliphatic glucosinolates (Met-derived) in both *mlk1/2/3* and *mlk1/3/4* mutant seedlings compared to wild type. In contrast, the levels of indolic glucosinolates (Trp-derived) remained unchanged in the mutant backgrounds (**Fig. 4**). Interestingly, the first seven glucosinolates originating from the earliest steps in the Met-derived glucosinolate biosynthetic pathway were increased 2-to 4-fold over wild type (**Fig. 4**), a pattern which correlates with the increased abundance of glucosinolate-associated biosynthetic enzymes (BCAT4, MAM1, SOT17/18, etc.) in the *mlk* mutants. Together these findings support a role for the MLKs in early stages of aliphatic glucosinolate biosynthesis and overall glucosinolate metabolism.

**Figure 4.**
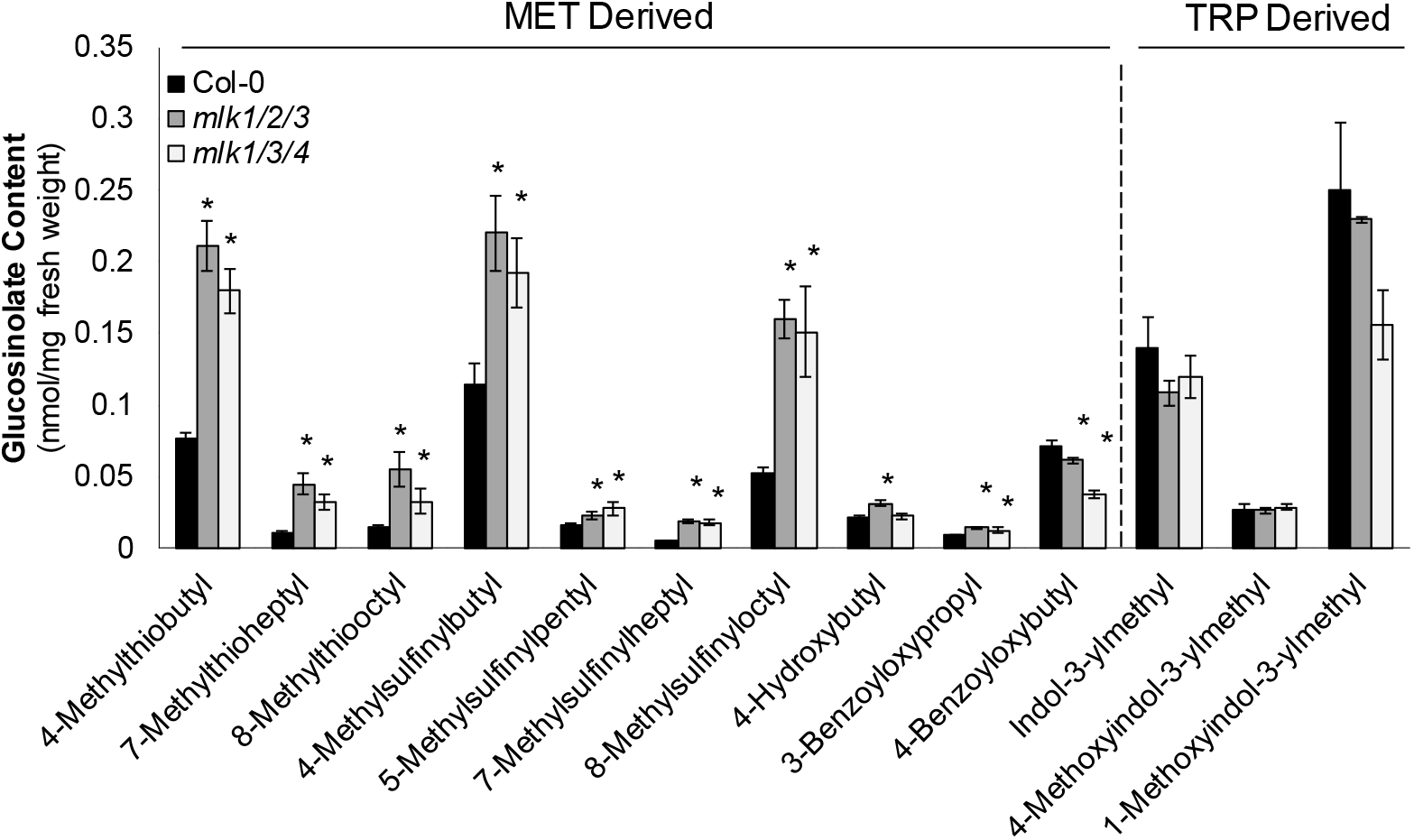
*mlk* Mutant Seedlings Contain Elevated Levels of MET Derived Glucosinolates. Glucosinolate (GLS) content of ten day old mutant and wild-type seedlings at ZT12 was quantified using HPLC. GLS identity determined using UV spectra and confirmed by LC-MS/MS. The average of four biological replicates is presented. Error bars indicate standard deviation. \**P*<0.01, compared with wild type seedlings (Student’s *t*-test).

In addition to glucosinolate biosynthetic enzymes, several proteins involved in hormone signaling and diverse stress responses changed in abundance in *mlk1/3/4* mutant seedlings compared to wild type at either ZT12 or ZT14, including BRI1-EMS-SUPPRESSOR 1 (BES1), SUPER SENSITIVE TO ABA AND DROUGHT2 (SAD2), CORONATINE INDUCED 1 (CORI3), pathogenesis-related gene 5 (PR5), lipoxygenase 2 (LOX2), thylakoidal ascorbate peroxidase (TAPX), and cold regulated 15a and b (COR15A and COR15B). Peptides mapping to the blue light receptor cryptochrome 2 (CRY2) also increased 2-fold in the *mlk1/3/4* mutant when compared to wild type at ZT12 (**Supplemental Dataset 1**). These observations are in agreement with the role of MLKs in hormone signaling, stress response, and light signaling (Liu et al., 2017; Dai and Xue, 2010; Casas-Mollano et al., 2008; Ni et al., 2017; Chen et al., 2018).

### Quantitative Phosphoproteomic Comparisons of *mlk* Mutants

The MLKs physically interact with and phosphorylate important regulatory proteins (Liu et al., 2017; Chen et al., 2018; Dai and Xue, 2010; Ni et al., 2017). Thus, we characterized the phosphoproteome of wild-type, *mlk1/2/3*, and *mlk1/3/4* mutant seedlings in the light (ZT12) and after transition to dark (ZT14) to gain insight into the role of the MLKs in global phosphorylation. We applied a TiO_2_ based phosphopeptide enrichment technique to the TMT-10plex labeled samples described above to achieve an in-depth phosphoproteomic analysis using the Thermo Scientific Orbitrap Fusion Lumos Tribrid mass spectrometer **(Fig. 1)**. Byonic software, run as a node within the Proteome Discover V2.1 platform identified and derived the relative quantitation of phosphoproteins. Using this strategy, we identified a combined total of 23,386 phosphosites on 15,222 unique peptides mapping to 4,854 protein groups at ZT12 and slightly fewer at ZT14 (19,947 phosphosites on 12,818 unique peptides mapping to 4,467 protein groups). At ZT12, over 80% of the identified phosphosites were serine residues, approximately 15% were threonine, and less than 2% were tyrosine (**Fig. 5A**), which is consistent with phosphosite distributions previously reported for Arabidopsis (Champion et al., 2004; Sugiyama et al., 2008). The phosphosite residue distributions were similar at ZT14 (**Figure 5A**).

**Figure 5.**
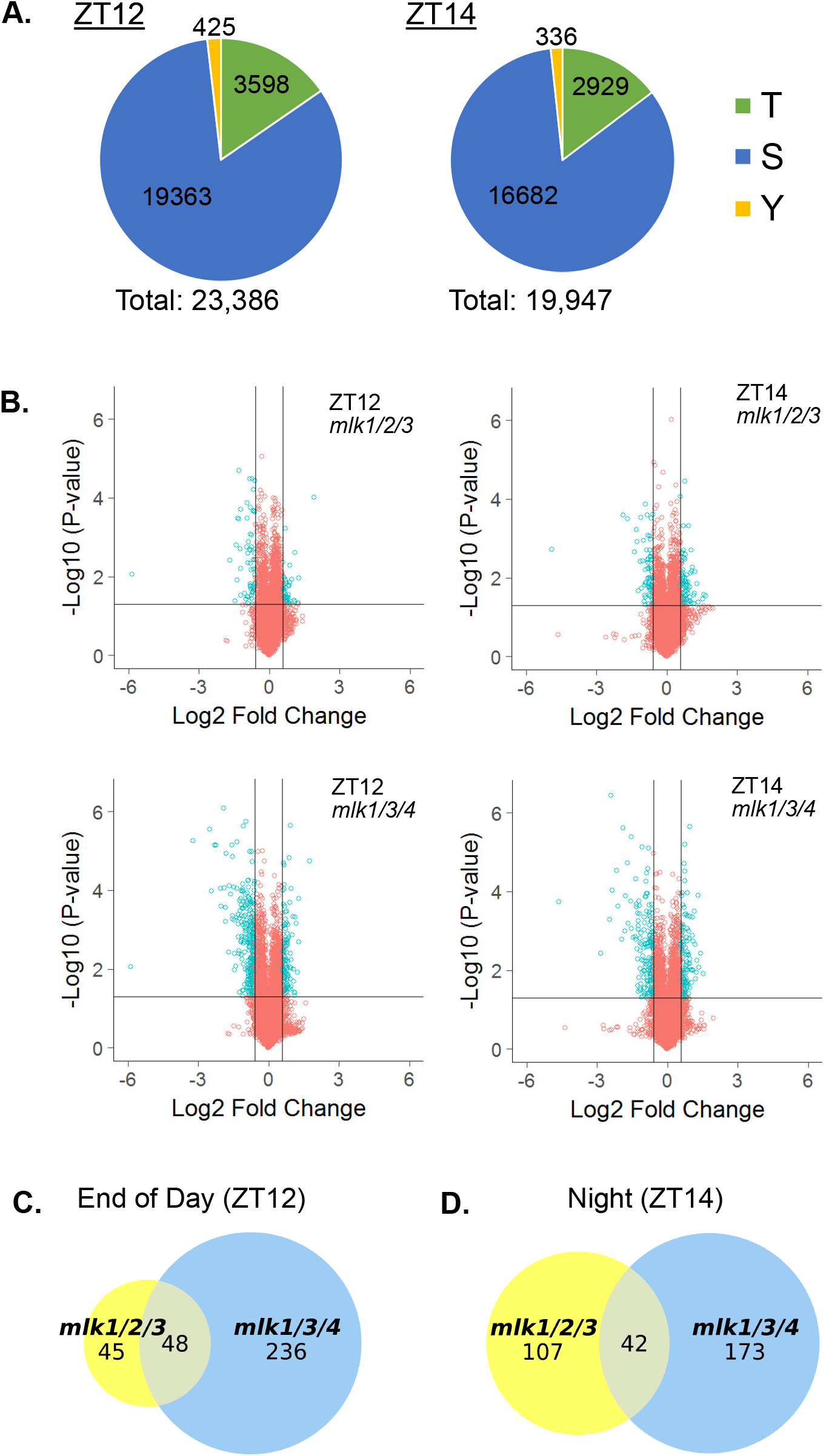
Analysis of Quantitative Phosphoproteomics of *mlk* Mutant Seedlings. (A) The distribution of threonine (T), serine (S), and tyrosine (Y) phosphorylation sites identified at ZT12 and ZT14. (B) Volcano plot of phosphopeptides identified in mutant and wild-type seedlings at ZT12 and ZT14. The x-axis specifies the log2 fold-change (FC) of mutant/wild-type and the y-axis specifies the negative logarithm to the base 10 of the t-test p-values. Open circles represent individual peptides, with blue circles specifying those considered statistically significant. Black vertical and horizontal lines reflect the filtering criteria (log2 FC = ±0.585 and p-value = 0.05) for significance. (C and D) Size-proportional Venn diagrams of differentially regulated phosphoproteins in *mlk1/2/3* and *mlk1/3/4* mutants at ZT12 (C) and ZT14 (D). Numbers indicate unique phosphoproteins.

We performed *mlk*-to-WT pairwise comparisons for each time point (**Supplemental Table S1 and Dataset S2**) to identify peptides that showed altered phosphorylation in the absence of the MLKs at either ZT12 or ZT14. We considered peptides differentially phosphorylated if they had a minimum log2 FC of ±0.585 (1.5 fold-change) and a p-value < 0.05. We identified 113 phosphopeptides corresponding to 93 unique protein groups that met the cutoff in the *mlk1/2/3* ZT12 set. In the *mlk1/3/4* ZT12 set, 429 phosphopeptides corresponding to 284 unique proteins were differentially phosphorylated, a 300% increase (**Fig. 5B**). 170 and 274 phosphopeptides, corresponding to 149 and 215 unique protein groups, have altered phosphorylation in the *mlk1/2/3* ZT14 and *mlk1/3/4* ZT14 analysis, respectively (**Figure. 5B and Supplemental Table S1**). The phosphosite residue distribution was similar in all datasets analyzed (**Figure 5A**). These data show that the *mlk1/3/4* mutant combination has a larger impact on the phosphoproteome than the *mlk1/2/3* mutant combination at each time point, particularly at ZT12. This observation supports a role for MLK4 in regulating the phosphoproteome in a light-dependent manner, likely through MLK4 phosphorylating blue light photoreceptors and red-light signaling pathway transcription factors (Liu et al., 2017; Ni et al., 2017).

Over 50% (48 out of 93) of the differentially phosphorylated proteins identified in the *mlk1/2/3* ZT12 set were present in the *mlk1/3/4* ZT12 set. In contrast, over 80% of those identified in the *mlk1/3/4* ZT12 set were specific to the *mlk1/3/4* mutant (**Fig. 5C**). When comparing the ZT14 sets, 42 phosphoproteins were shared, accounting for 28% and 19.5% of the proteins identified in the *mlk1/2/3* ZT14 and *mlk1/3/4* ZT14 sets, respectively (**Fig. 5D**). Many of the proteins shared between *mlk1/2/3* and *mlk1/3/4* protein sets are involved in gene silencing and chromatin organization (**Supplemental Dataset S2**), supporting a conserved role for the MLK family kinases in these processes (Casas-Mollano et al., 2008; Jeong Br et al., 2002). Some of these proteins, including SUO and SERRATE (SE) – which are involved in microRNA biogenesis pathways, exhibit altered phosphopeptide abundance at both ZT12 and ZT14. However, proteins involved in chromatin organization, such as Increased in Bonsai Methylation 1 (IBM1) and SPLAYED (SYD), only showed altered phosphorylation in both *mlk1/2/3* and *mlk1/3/4* mutant backgrounds at ZT12 (**Supplemental Dataset S2**). These results suggest that the MLKs are involved in regulating gene expression, possibly through modulating light-dependent chromatin organization.

### MLKs Influence Diverse Kinase Signaling Networks

The proteins identified as differentially phosphorylated in the *mlk1/3/4* mutant background at both ZT12 and ZT14 are associated with a diverse set of biological processes, suggesting a possible disruption of multiple protein kinase networks. Motif-X (http://motif-x.med.harvard.edu/motif-x.html; (Chou and Schwartz, 2011; Schwartz and Gygi, 2005) was used to isolate overrepresented sequence motifs present in the phosphopeptide sets that are associated with known kinase families. Following extension of differentially phosphorylated high-confidence, unambiguous peptides using PEPTIDEXTENDER ver.0.2.2 alpha (http://schwartzlab.uconn.edu/pepextend), the resulting 15-mers were submitted for motif analysis using a significance threshold of p < 10^−6^ and a minimum occurrence requirement of 20. Peptides exhibiting increased or decreased abundance when compared to wild type were analyzed separately. Three serine phosphorylation (Sp) motifs (S-x-x-K, R-x-x-S, and K-x-x-S) were overrepresented in peptides that were decreased in abundance at ZT12 in the *mlk1/3/4* mutant seedling background (**Fig. 6A**). The K-x-x-S motif, along with an acidic S-type motif (S-x-x-x-x-x-E), was also overrepresented at ZT14 in *mlk1/3/4* mutants (**Fig. 6B**). The CDPK-SnRK superfamily of protein kinases is known to recognize R/K-x-x-S/T basic motifs. The R-x-x-S motif has also been associated with the AGC family kinases, PKA and PKC (Rademacher and Offringa, 2012; Marondedze et al., 2016), which are involved in mid-to late-day rhythmic phosphorylation (Choudhary et al., 2015). The kinase(s) responsible for phosphorylation at the S-x-x-K site in plants is unknown. However, the highly conserved eukaryotic cyclin B-dependent protein kinase Cdk1 that recognizes several non-S/T-P motifs, including the S/T-x-x-R/K motif, is a candidate (Suzuki et al., 2015). The classical minimal motif required for recognition by proline-directed kinases families (mitogen-activated protein kinase (MAPK), cyclin-dependent kinase (CDK), and glycogen synthase kinase 3 (GSK-3)), S-P, was found to be overrepresented in peptides that were increased in abundance in *mlk1/3/4* mutants at ZT12 and ZT14. While the S-x-S motif, which is associated with the receptor-like protein kinase (RLK) family (van Wijk et al., 2014), was only found to be overrepresented at ZT12 (**Fig. 6**). Using these parameters, there were no overrepresented motifs identified from the sites that decreased in abundance in either of the *mlk1/2/3* data sets. However, if we reduced the minimum occurrence to 10, then the R-x-x-S and S-P motifs were overrepresented in peptides with decreased or increased abundance at ZT12 in *mlk1/2/3* mutants (**Supplemental Figure S2**). The diversity of identified overrepresented kinases motifs suggests that MLK family kinases influence numerous biological processes through systemic regulation of multiple kinase signaling networks.

**Figure 6.**
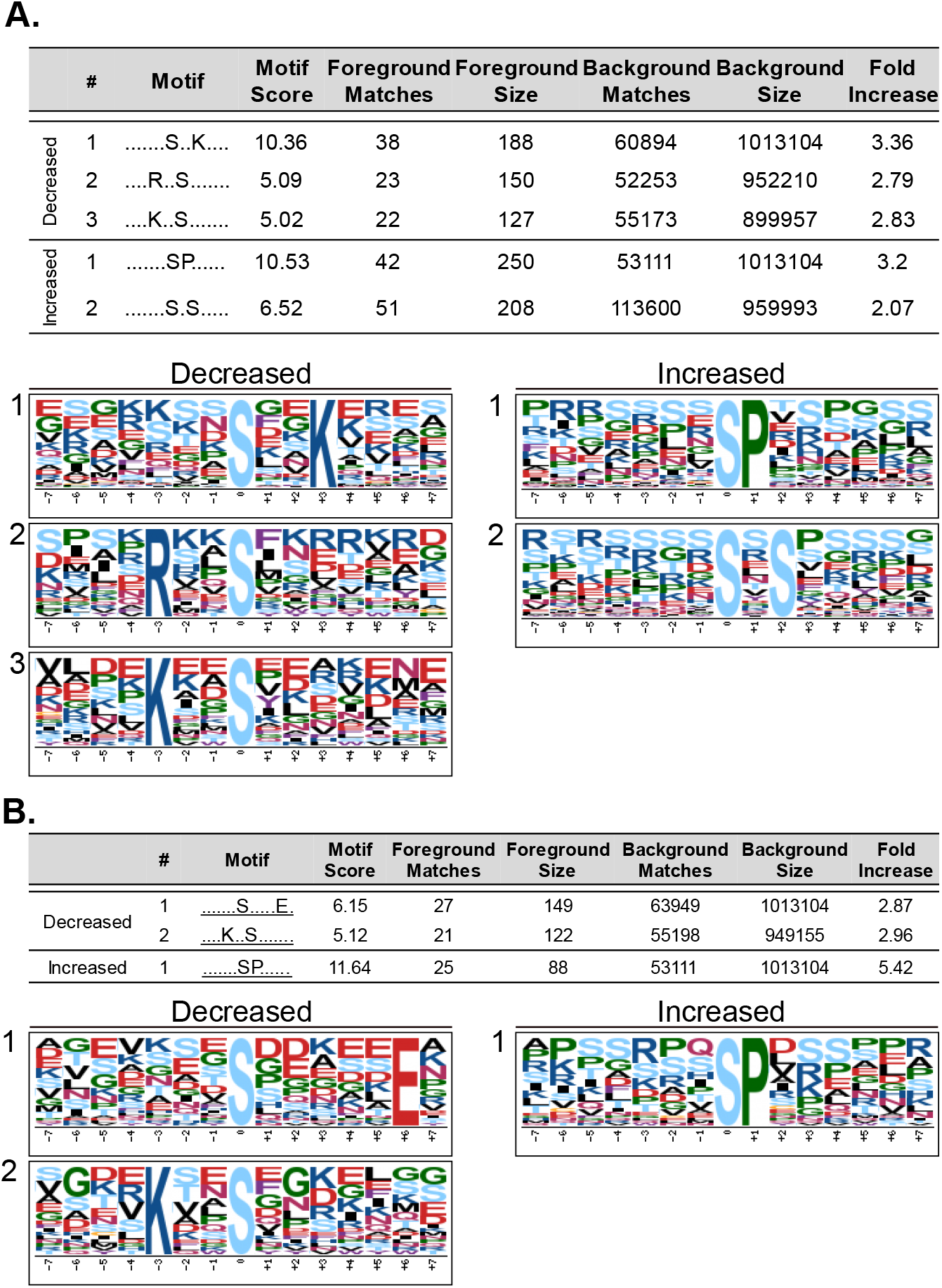
Motif Analysis of Differentially Phosphorylated Peptides. Phosphopeptides with altered abundance in *mlk1/3/4* mutants at ZT12 (A) and ZT14 (B) where extended (http://schwartzlab.uconn.edu/pepextend) and centered. Motif-X analysis was then preformed with the probability threshold was set to p-value ≤ 10^−6^, the occurrence threshold was set to 20, and the default IPI Arabidopsis Proteome data set was used as the background data set.

### MLKs Influence the Phosphorylation Status of Nuclear Localized Proteins

Using g:Profiler (https://biit.cs.ut.ee/gprofiler/), 149 GO terms were enriched in the differentially phosphorylated proteins from the *mlk1/3/4 Z*T12 set; 107 of these were classified as biological process, 34 as cellular component, and 5 as molecular function. Forty-nine terms were enriched in the *mlk1/3/4 Z*T14 set, 28 in biological process and 21 in cellular component. Fewer terms were found to be enriched in the *mlk1/2/3* phosphoprotein sets, with only 8 enriched terms at ZT12 and 18 at ZT14. A complete list of enriched GO terms can be found in **Supplemental Dataset S2**. Functionally redundant terms were removed using REVIGO (default settings, dispensability threshold = 0.7 (cellular component) or 0.5 (biological process); **Supplemental Tables S2-3;** (Supek et al., 2011)). Under the cellular component category, there was strong enrichment for terms associated with the nucleus: ‘nucleus’ (GO:0005634), ‘nuclear part’ (GO:0044428), and ‘nucleoplasm’ (GO:0005654). Terms including ‘chromosome’ (GO:0005694), ‘chromosomal part’ (GO:0044427), and ‘chromatin’ (GO:0000785) were also found to be enriched in at least one of the differentially phosphorylated protein lists (**Fig. 7A**). These results are in agreement with the known nuclear localization of the MLKs, and their role in modifying chromatin (Wang et al., 2015a; Huang et al., 2016; Su et al., 2017).

**Figure 7.**
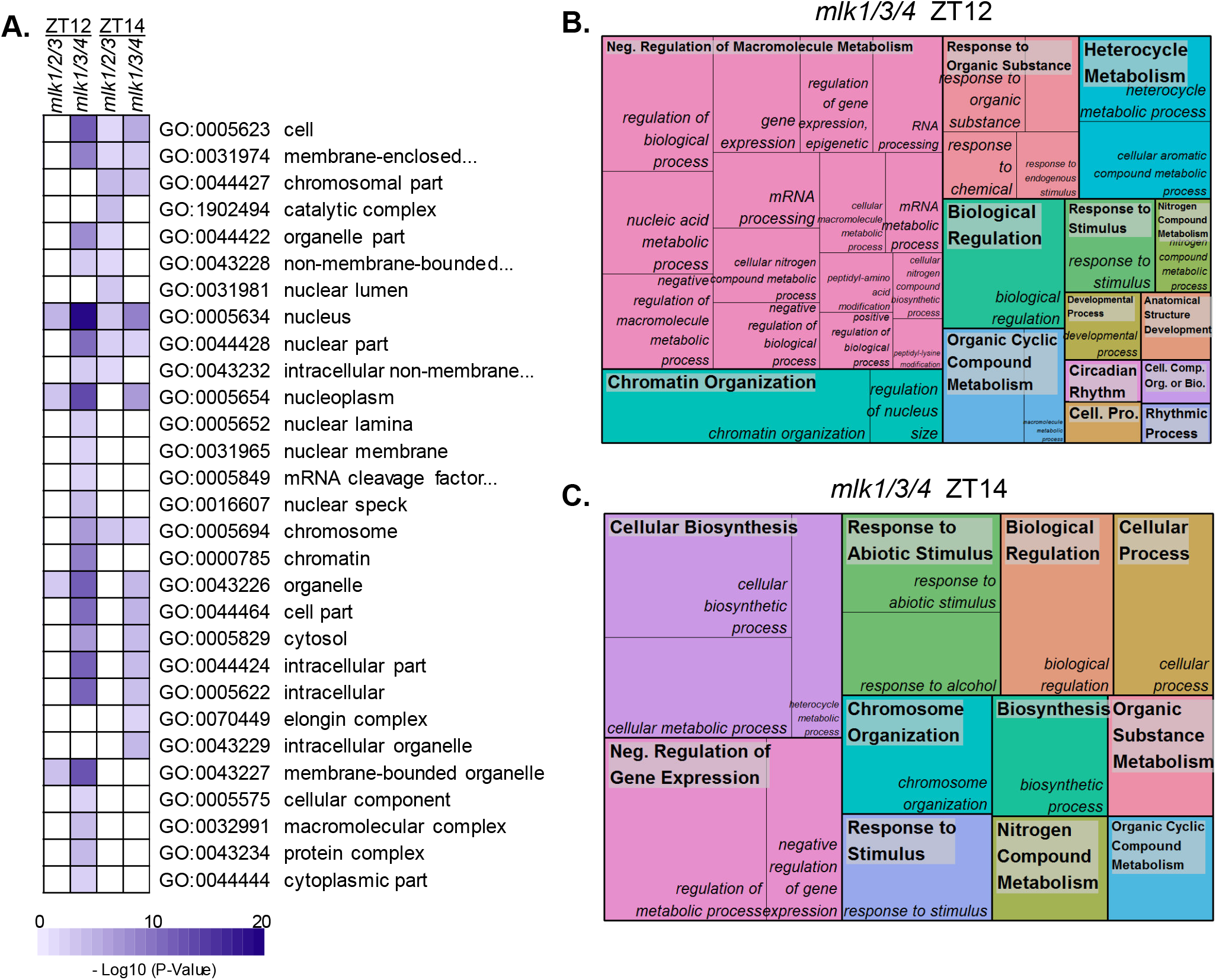
Gene Ontology Enrichment Analysis of Differentially Phosphorylated Proteins. (A) Heat map showing the p-value significance of enriched cellular component GO categories of proteins with altered abundance in *mlk* mutant seedlings at ZT12 and ZT14. (B and C) Treemap representation of Biological Process GO categories enriched in *mlk1/3/4* mutant seedlings at ZT12 (B) and ZT14 (C). The box size correlates to the −log10 *P*-value of the GO-term enrichment. Boxes with the same color indicate related GO-terms and correspond to the representative GO-term which is found in the upper-left of each box. REVIGO was used to eliminate redundant GO-terms with a dispensability value ≥ 0.7 (A) or ≥ 0.5 (B and C).

### Circadian-Associated Proteins Exhibit Altered Phosphorylation in *mlk1/3/4* Mutant Seedlings

The *mlk1/3/4* ZT12 differentially phosphorylated protein set was enriched in proteins associated with rhythmic processes (GO:0048511) and/or circadian rhythms (GO:0007623). These observations agree with previous reports linking the MLKs to light signaling and circadian regulation (Huang et al., 2016; Ni et al., 2017; Liu et al., 2017; Su et al., 2017; Zheng et al., 2017). The core circadian clock proteins PSEUDO-RESPONSE REGULATOR 7 (PRR7), TIME FOR COFFEE (TIC), and REVEILLE 8 (RVE8) were all differentially phosphorylated in the *mlk1/3/4* mutant background at ZT12 (**Table 1 and Supplemental Table S3**). We observed decreased phosphorylation of TIC at S324 and PRR7 at S355 and S275, while RVE8 showed increased phosphorylation of the C-terminal half (**Table 1**). The red-light photoreceptor phytochrome B (PHYB) and the transcriptional master regulator ELONGATED HYPOCOTYL 5 (HY5) also showed reduced phosphorylation at Threonine 42 and T64, respectively (**Table 1**); to the best of our knowledge, these phosphosites are previously unreported. **Table 1** lists additional circadian-associated proteins with altered phospho-abundance in the *mlk1/3/4* mutant at ZT12.

**Table 1.**
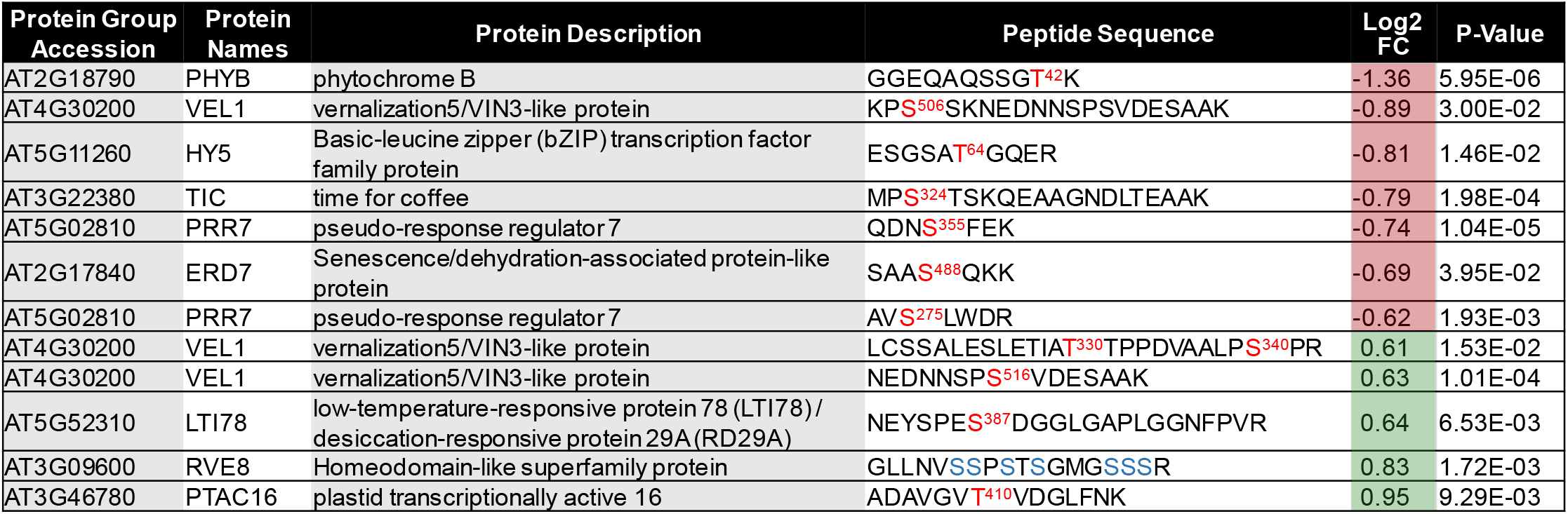
Phosphosites Identified in Circadian-Associated Proteins.

### Gene Ontology Analysis Reveals Enrichment of Proteins Involved in Chromatin Organization

Due to the large number of enriched GO terms identified from both the *mlk1/3/4 Z*T12 and *mlk1/3/4 Z*T14 list, the dispensability threshold was reduced to 0.5 for REVIGO analysis. This resulted in the identification of 34 and 15 representative and non-redundant enriched biological process terms in *mlk1/3/4* ZT12 and *mlk1/3/4 Z*T14 altered phosphoprotein lists, respectively. Enriched GO terms and their underlying gene identifiers shared between the *mlk1/3/4 Z*T12 and ZT14 sets included, ‘organic cyclic compound metabolism’, GO: 1901360, ‘nitrogen compound metabolism’, GO:0006807, ‘chromosome organization’, GO:0051276 and ‘negative regulation of gene expression’ GO:0006807 (**Fig. 7B-C and Supplemental Table S3**). The chromatin modifying proteins BRAHMA (BRM), SIN3-like3 (SNL3), vernalization5/VIN3-like (VEL1), and high mobility ground B1 (HMGB1) were shared amongst these GO terms. Additional proteins associated with chromatin modifications were present in the *mlk1/3/4 Z*T12 list including the histone methyltransferase EARLY FLOWERING IN SHORT DAYS (EFS)/SET DOMAIN GROUP 8 (SDG8), the histone acetyltransferase TBP-ASSOCIATED FACTOR 1 (TAF1), as well as IBM1, actin-related protein 4 (ARP4), alfin-like 7 (AL7), GLIOMAS 41 (GAS41/YAF9a), and stress-induced histone H2A protein 9 (HTA9). Few biological process GO-terms were found to be enriched in the *mlk1/2/3* data sets. Nevertheless, those that were (‘regulation of gene expression’, ‘epigenetic’ (ZT12) and ‘chromosome organization’, ‘chromatin organization’ and ‘mitotic sister chromatid cohesion’ (ZT14)) were also enriched in the *mlk1/3/4* protein list (**Supplemental Dataset S2**). These results support a role for the MLKs in regulating chromatin organization and gene expression at the assessed time points.

### *mlk1/3/4* Mutants Show Altered Phosphorylation of Proteins Involved in Nuclear Organization and DNA Damage Response

Analysis of differentially phosphorylated peptides has shown that the loss of *mlk1, mlk3*, and *mlk4* at ZT12 has the greatest impact on both the global- and phosphoproteome. Therefore, we chose to expand on our analysis exclusively for the *mlk1/3/4* ZT12 data set. To elucidate further the biological processes influenced by the MLKs at the end of the day (ZT12), we independently analyzed the phosphoproteins that were increased or decreased in abundance. Of the 429 phosphopeptides found to have altered abundance in the *mlk1/3/4* mutant background at ZT12 (**Supplemental Dataset S2**), 133 were increased and 296 were decreased in abundance, mapping to 103 increased and 190 decreased unique proteins. Interestingly, 9 gene identifiers were shared between the increased and decreased groups, including several that are involved in various aspects of nuclear organization such as LITTLE NUCLEI 1/ CROWED NUCLEI 1 (LINC1/CRWN1), VEL1, and HMGB1 (**Fig. 8A and Supplemental Dataset 2**). Proteins that were increased in abundance were associated with the representative GO-terms ‘response to organic substance’ (GO:0010033) and ‘response to stimulus’ (GO:0050896) (**Fig. 8B**). However, we found that the majority of GO terms enriched in the inclusive set (both increased and decreased peptides) such as ‘RNA processing’ (GO:0006396), ‘chromosome organization’ (GO:0051276), and ‘developmental processes’ (GO:0032592) are associated with decreased phosphorylation (**Fig. 8C**). The GO term ‘cellular response to DNA damage stimulus’ (GO:0006974), was also enriched in proteins exhibiting decreased phosphorylation. Proteins associated with this term include a catalytic subunit of DNA polymerase alpha INCARVATA2 (ICU2), as well as X-ray cross complementation group4 (XRCC4) and MUTM homolog-1 (MMH-1) both of which are directly involved in DNA repair (West et al., 2000; Ohtsubo et al., 1998; Barrero et al., 2007).

**Figure 8.**
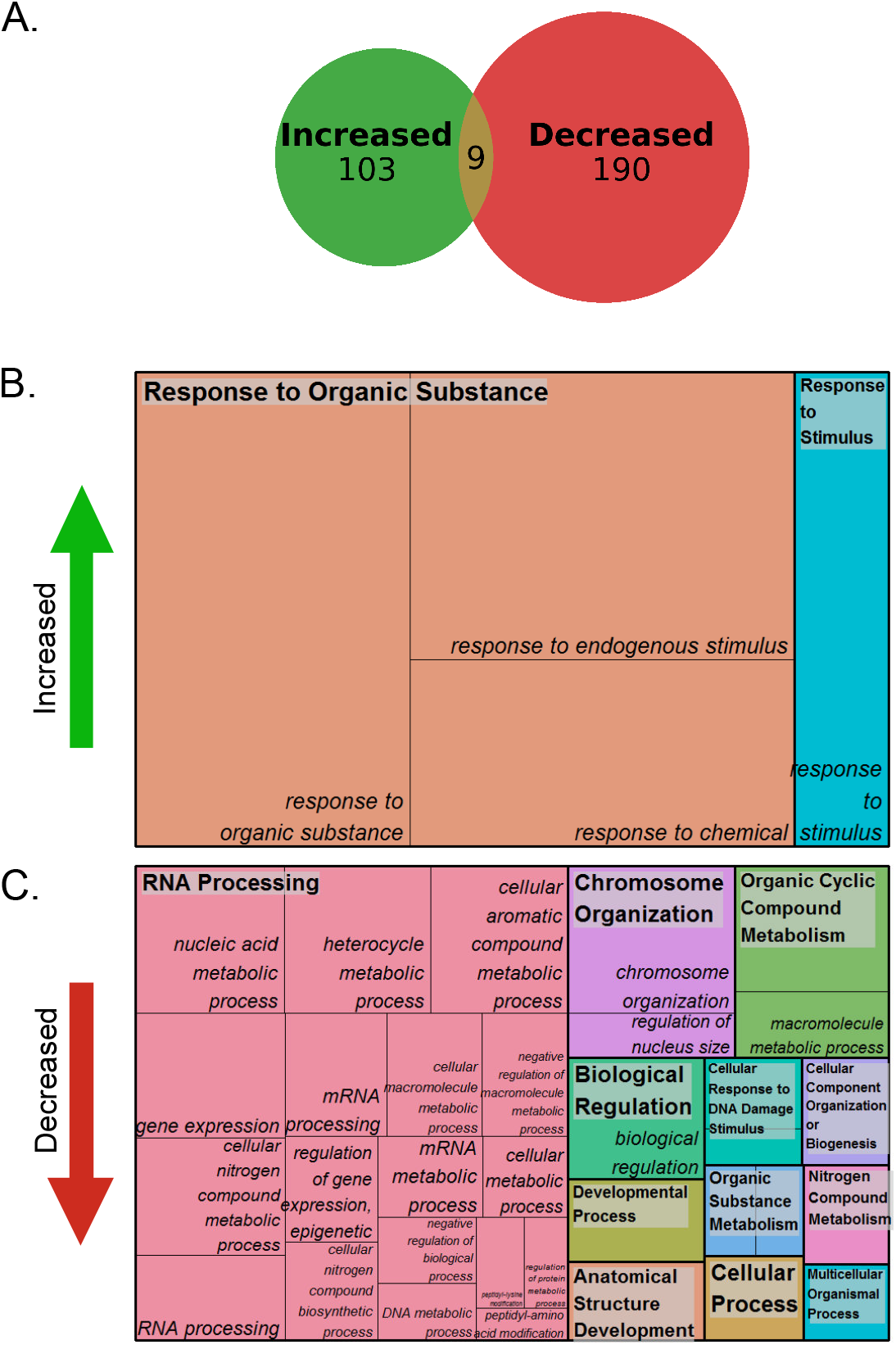
Analysis of Differentially Phosphorylated Peptides in *mlk1/3/4* Mutants at ZT12. (A) Size-proportional Venn diagram of proteins which show increased, decreased, or both increased and decreased abundance of identified phosphosites in *mlk1/3/4* mutant seedlings at ZT12. Numbers indicate unique phosphoproteins. Analysis of phosphoproteins that show increased (B) or decreased (C) in abundance in the *mlk1/3/4* mutant seedlings background at ZT12. Treemap representations of Biological Process GO category enrichment are shown. The box size correlates to the −log10 *P*-value of the GO-term enrichment. Boxes with the same color indicate related GO-terms and correspond to the representative GO-term which is found in the upper-left of each box. REVIGO was used to eliminate redundant GO-terms with a dispensability value ≥ 0.5.

### *mlk* Mutants Show Increased Sensitivity to DNA-damaging Agents

Nuclear organization and chromatin dynamics strongly influence DNA damage repair efficacy (Reviewed in Vergara and Gutierrez, 2017; Donà and Mittelsten Scheid, 2015). Additionally, the *Chlamydomonas* MLK orthologue, Mut9, is required for survival when grown in the presence of genotoxic agents (Jeong Br et al., 2002). Since many of the proteins exhibiting changes in phosphorylation abundance in the *mlk1/3/4* mutants at ZT12 are proteins associated with nuclear organization (LINC/CRWN family members, SAD1/UNC-84 domain protein 2 (SUN2), BRM, and SYD) and DNA damage, we further explored what role the *Arabidopsis* MLKs might play in DNA damage response. To do so, we evaluated the sensitivity of mutant and wild-type seedlings to the genotoxic agent methyl methane sulfonate (MMS) and UV-C. MMS is a monofunctional DNA alkylating agent that induces replication fork stalling and subsequent double strand breaks (Ensminger et al., 2014). In addition to the *mlk1/2/3* and *mlk1/3/4* mutants, the *mlk4* single and *mlk* quadruple amiRNA line (amiR^4k^)(Liu et al., 2017) were included in our analysis.

Mutant and wild-type seedlings were imaged and weighed after fourteen days of exposure to titrations of MMS. All genotypes showed reduced aerial mass with increasing levels of MMS, with the *mlk* mutants showing increased sensitivity compared to wild type (**Fig. 9A-B**). In wild-type seedlings, growth was reduced by less than 10% (fresh weight) in the presence of 50 ppm MMS compared to the mock-treated seedlings. The *mlk4* single mutant seedlings showed more than 20% reduction of fresh weight, the *mlk1/3/4* seedlings a 39%, and amiR^4K^ lines a 44% reduction, compared to the mock-treated seedlings (**Fig. 9B**). The *mlk1/3/4* and amiR^4k^ mutants continued to decline in fresh weight at higher concentrations of MMS. At 100 ppm MMS, *mlk1/2/3* seedlings showed approximately a 10% greater reduction in fresh weight than what was observed for wild type (**Fig. 9B**). In addition to stunted growth, chlorotic tissue was observed in the *mlk1/3/4* mutants growing on 50 ppm MMS and in the amiR^4K^ mutants at 75 PPM MMS (**Fig. 9A**). 150 ppm MMS caused severe growth reduction and lethality in all assessed genotypes; thus, seedlings were imaged but not weighed (**Fig. 9A**). Next, we assessed seedling germination and growth in the continuous presence of 150 ppm MMS. Post-germination growth was severely impaired in the *mlk1/3/4* mutant, with more than 65% of seedlings exhibiting complete developmental arrest compared to approximately 10% of wild-type seedlings (**Fig. 9C-D**). In contrast, the *amiR^4k^* mutant seedlings developed similar to wild-type seedlings when germinated in the presence of 150 ppm MMS (**Fig. 9C-D**), which could be a result of near endogenous expression levels of MLK2 and MLK3 (Ni 2017).

**Figure 9.**
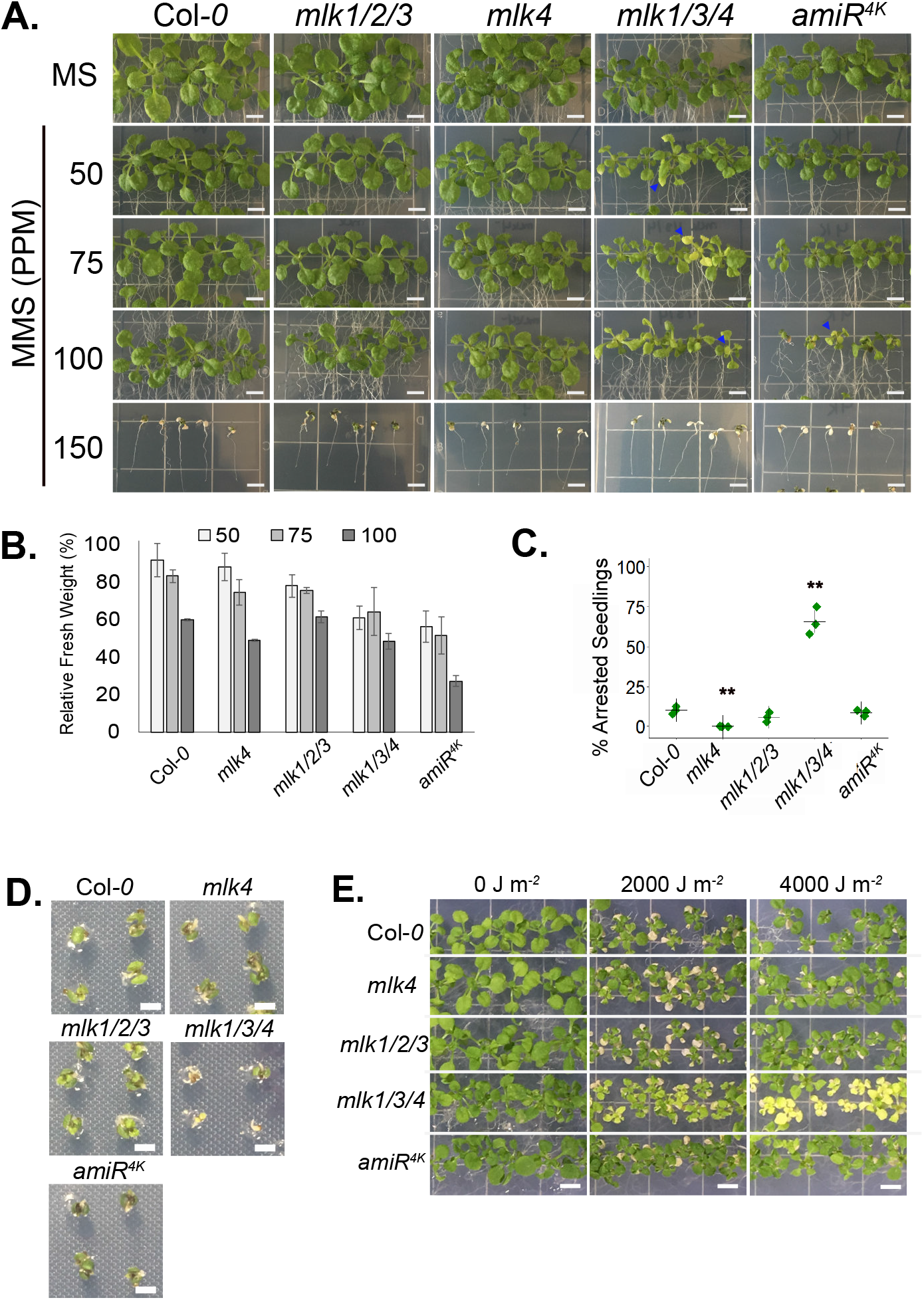
*mlk1/3/4* Mutants Have Increased Sensitivity to MMS Treatment. (A) Representative images of mutant and wild-type seedlings 15 days after transfer to solid media containing the indicated concentration of MMS. Chlorotic tissue is denoted with a blue arrowhead. Scale bar = 5 mm. (B) Fresh weight of 21-day old wild type and mutant seedlings grown in the presence of MMS relative to mock treated samples. The average of 3 biological replicates of ≥10 seedlings each is presented. (C) Percent of seedlings exhibiting post-germination developmental arrest after 12 days of growth in the presence of 150 PMM MMS. The average of 3 biological replicates of ≥30 seedlings is presented. (B and C) Error bars indicate standard deviation. \**P*<0.05, \*\**P*<0.01compared with wild-type seedlings (Student’s *t*-test). (D) Representative images of mutant and wild type seedlings germinated in the presence of 150 PPM MMS. (E) Representative images of mutant and wild-type seedlings irradiated with indicated levels of UV-C. Scale bar = 2 mm.

We also tested the *mlk* mutants for sensitivity to UV-induced DNA damage through periodic exposure to multiple doses of UV-C irradiation. The phenotypic impact of chronic irradiation with either 2000 or 4000 J m^−2^ on seedlings was evaluated five days after a recovery period. All genotypes had cotyledon cell death and reduced growth after exposure to both doses of UV-C. However, only *mlk1/3/4* mutant seedlings showed tissue chlorosis after exposure to both 2000 and 4000 J m^−2^ UV-C. amiR^4k^ mutants had minimal chlorosis after irradiation with 4000 J m^−2^ (**Fig. 9E**). Taken together, these data suggest that *mlk* mutants have increased sensitivity to DNA damage.

## Discussion

### MLK protein kinases alter protein phosphorylation in developmental and stress responsive pathways

The current repertoire of MLK substrates is composed of a photoreceptor, multiple transcription factors, hormone receptors, and histones (Ni et al., 2017; Liu et al., 2017; Dai and Xue, 2010; Chen et al., 2018; Casas-Mollano et al., 2008; Su et al., 2017; Kang et al., 2020; Wang et al., 2015a). MLK mutants have defects in circadian period length, hypocotyl elongation, flowering time, osmotic stress responses, seed set, chromatin organization, and hormone sensitivity (Su et al., 2017; Casas-Mollano et al., 2008; Liu et al., 2017; Huang et al., 2016; Zheng et al., 2017; Chen et al., 2018; Kang et al., 2020). These observations support a model where MLK family members function as central regulators of numerous interconnected signaling pathways. Our quantitative analysis of the global- and phosphoproteomes of *mlk* triple mutant seedlings supports a diverse and complex role for the MLKs in the regulation of cellular signaling and response. Interestingly, these kinases seem to share a balance of functional redundancy and substrate specificity, which for example, results in opposing circadian period and hypocotyl elongation phenotypes (Huang et al., 2016; Liu et al., 2017). We found the *mlk1/3/4* mutant displays a much more severe proteomic phenotype relative to the *mlk1/2/3* mutant, with over 10-fold more proteins showing altered abundance in the *mlk1/3/4* mutant (**Fig. 2**). This increase holds for the phosphoproteome as well. However, the difference was greatest in tissue sampled in the light. An explanation for the light dependence could be the result of MLK4 acquiring substrate specificity or MLK4 having a greater tendency than other MLKs for interacting with light-signaling proteins in planta. MLK4 has a higher affinity for PIF3 when compared to other MLKs (Ni et al., 2017), and both phyB and HY5 have altered phosphorylation only at ZT12 in the *mlk1/3/4* mutants (**Table 1**). While some proteins with altered abundance were unique to the *mlk1/3/4* or *mlk1/2/3* mutants, shared targets include glucosinolate biosynthesis (global proteome; **Fig. 3**) and chromosome organization (phosphoproteome; **Fig. 7B-C and Supplemental Dataset S2**). Further work is needed to determine if any of the proteins showing altered phosphorylation are direct substrates of the MLKs, or whether changes in phosphorylation status is occurring indirectly through additional kinases.

### MLKs Regulate Hormone Signaling and Stress Responses

Several proteins responsible for glucosinolate metabolism showed altered abundance in *mlk* mutant seedlings before and after dark transition (**Fig. 3** and **Supplemental Dataset S1**). Glucosinolates are nitrogen- and sulfur-containing secondary metabolites known for their role in plant defense (Kim et al., 2008; Kos et al., 2012; Bednarek et al., 2009; Clay et al., 2009) and anticarcinogenic properties (Higdon et al., 2007). Accumulation of glucosinolates in *Arabidopsis thaliana* is rhythmic, controlled in part by circadian clock regulated jasmonate accumulation and the activity of the basic leucine zipper (bZIP) transcription factor, ELONGATED HYPOCOTYL 5 (HY5) (Goodspeed et al., 2012; Huseby et al., 2013). HY5 is a positive regulator of light signaling and functions as a central regulator of light-dependent growth and development by integrating various environmental signals (Gangappa and Botto, 2016). Peak glucosinolate levels occur during the day, possibly to protect against rhythmic herbivory. Two glucosinolate biosynthesis genes that showed increased protein abundance in *mlk* mutants, *CYP79F1* and *SOT18*, are expressed at lower levels in the *hy5* mutant background (Huseby et al., 2013). The phosphorylation of HY5 is associated with its activity and stability, with the non-phosphorylated form being more active (Hardtke, 2000). Thus, it is possible that increased HY5 activity, resulting from decreased phosphorylation in the *mlk* mutant background, could be influencing glucosinolate metabolism. Additionally, abscisic acid (ABA) induces glucosinolate accumulation in plants (Wang et al., 2015b; Zhu and Assmann, 2017). MLK3 regulates ABA signaling through the phosphorylation of the PYR/PYL ABA receptor family of proteins (Chen et al., 2018). In agreement with altered ABA signaling, increased phosphorylation of proteins associated with ABA responses and the SnRK consensus motif, R/K-x-x-S/T, was found to overrepresented in the *mlk1/3/4* mutant background. Thus, the MLKs may be involved in the regulation of defense responses through multiple converging signaling pathways.

### The Phosphorylation Status of Key Circadian and Light Signaling Components Are Altered in the Absence of MLK Family Kinases

Several differentially phosphorylated proteins that are involved in chromatin organization also function as core circadian clock components (RVE8) or are central regulators of clock input pathways such as temperature and light signaling (phyB and HY5). RVE8 is a MYB-like transcription factor that regulates the expression of the clock gene *TIMING OF CAB EXPRESSION1 (TOC1*) by promoting histone 3 (H3) acetylation of its promoter (Farinas and Mas, 2011). RVE8 shares structural similarity to the core clock transcription factors CCA1 and LATE ELONGATED HYPOCOTYL (LHY). Phosphorylation of CCA1 by the Ser/Thr protein kinase CK2 antagonistically regulates CCA1 transcriptional activity by reducing its ability to bind to the promoters of clock gene targets, which in turn alters circadian period (Portolés and Más, 2010). Further exploration of the impact of RVE8 phosphorylation could reveal a new avenue of post-translation regulation of the circadian clock.

Temperature and light signaling are critical circadian inputs that allow plants to coordinate growth and development (*e.g*., germination and photoperiodic flowering) with their environment. The phyB photoreceptor is central to both temperature and light signaling pathways (Legris et al., 2016). PhyB activity is regulated in part by phosphorylation of its N-terminus (Medzihradszky et al., 2013; Nito et al., 2013). Altered phospho-status of phyB Ser86 and Y104 influences phyB rate dark-reversion rates, hypocotyl elongation, and flowering time in Arabidopsis (Hajdu et al., 2015; Medzihradszky et al., 2013; Nito et al., 2013). Here we report decreased phosphorylation of a previously unidentified phyB N-terminal phosphosite, T42, in the *mlk* mutant background at ZT12 (**Table 1**). MLKs are known to associate with phyB, phosphorylate the phytochrome–interacting factor, PIF3, and display a variety of red-light dependent growth phenotypes (Ni et al., 2017; Huang et al., 2016). In addition to the well-established light-induced phyB-PIF signaling cascade, there is an ample amount of evidence supporting the role of phyB in large-scale chromatin organization (van Zanten et al., 2010; Tessadori et al., 2009). Thus, the MLK-phyB interaction may contribute to light-dependent chromatin re-organization in addition to regulating PIF3 turnover. We also found decreased phosphorylation of another key light signaling component, HY5 at T64. Whether the phosphorylation of HY5^T64^ and/or phyB^T42^ is directly or indirectly influenced by the MLKs and how those phosphosites fit into the existing light signaling paradigm will be an exciting line of future research.

### The Role of MLK Family Kinases in Modulating Nuclear Architecture

The role of histone modifications in the regulation of developmental processes and stress response has been well-established, yet our understanding of the responsible modifiers, modification crosstalk, and targeted genes is incomplete (Probst and Mittelsten Scheid, 2015; Rosa and Shaw, 2013). Early observations have implicated the MLK family in the regulation of environmentally-stimulated chromatin organization. MLK1, like its *Chlamydomonas* homologue MUT9, has been shown to phosphorylate histone H3 on threonine 3 (H3T3p) and to function redundantly with MLK2 to promote H3T3p in response to salt stress (Wang et al., 2015a; Casas-Mollano et al., 2008). Accordingly, the *mlk1mlk2* double mutant has abnormal chromatin organization and increased sensitivity to osmotic stress (Wang et al., 2015a). Comparisons have been drawn between the defects in chromosomal organization observed in the *mlk1mlk2* mutants and those occurring in plants harboring mutations in members of the LITTLE NUCLEI/CROWDED NUCLEI (LINC/CRWN) gene family, which are involved in controlling nuclear size and heterochromatin organization (Wang et al., 2013; Sakamoto and Takagi, 2013). Our analysis of the *mlk* mutant phosphoproteomes found that peptides mapping to multiple members of the LINC/CRWN family were altered in abundance in *mlk1/2/3* and *mlk1/3/4* mutants, suggesting that MLKs may influence nuclear organization in part through regulation of the LINC proteins.

### MLKs Are Involved in DNA Damage Repair Through Multiple Pathways

Plants are exposed to DNA-damage from their external environment (e.g., ultraviolet light, ionizing radiation, heat stress, and bacterial and fungal toxins) as well as endogenous sources such as DNA-alkylating metabolic byproducts. Maintenance of genomic integrity requires an efficient DNA damage repair (DDR) system that can identify, access, remove, and reassemble damaged genomic regions within the context of chromatin. Mutations in genes involved in chromatin organization and remodeling often exhibit defects in DDR and enhanced susceptibility to DNA damaging reagents (Donà and Mittelsten Scheid, 2015). The increased sensitivity of *mlk* mutants to DNA damaging agents could result from the dysregulation of proteins involved in chromatin remodeling, such as GAS41/YAFa, ARP4, BRM, and SYD (**Fig. 9**). The *mlk1/3/4* mutant also shows altered phosphorylation of several proteins directly involved in DDR, such as MMH-1 and XRCC4 (Yuan et al., 2014; Roy et al., 2013). Additionally, there is accumulating evidence supporting a role for small regulatory RNAs in DDR (Hawley et al., 2017); proteins associated with small RNA metabolism are enriched in *mlk1/3/4* at ZT12 (**Fig. 7**). There is no question that full elucidation and validation of the mechanisms linking MLKs and DDR will require further exploration. However, taken together, our data suggests the MLKs play an important role in mitigating DNA damage through the regulation of multiple response pathways.

## Supporting information

Supplemental Data set S1

Supplemental Data Set S2

