## Supplemental Figures for "Quantitative Proteomics and Phosphoproteomics Supports a Role for Mut9-Like Kinases in Multiple Metabolic and Signaling Pathways in *Arabidopsis*"

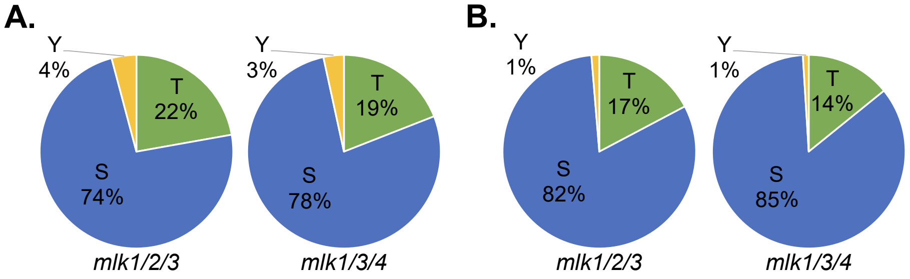


**Supplemental Figure S1. Phosphosite distribution of differentially phosphorylated peptides in *mlk* mutants**

The of threonine (T) phosphorylation, serine (S) phosphorylation, and tyrosine (Y) phosphorylation sites identified as having altered abundance in *mlk1/2/3* or *mlk1/3/4* mutants compared to wildtype samples at ZT12 (A) or ZT14 (B).

**A.**

| | # | Motif | Motif Score | Foreground Matches | Foreground Size | Background Matches | Background Size | Fold Increase |
|---|---|---|---|---|---|---|---|---|
| Decreased | 1 | ....R..S....... | 5.08 | 12 | 48 | 55686 | 1013104 | 4.55 |
| Increased | 1 | .......SP...... | 5.71 | 11 | 37 | 53111 | 1013104 | 5.67 |

**B.**

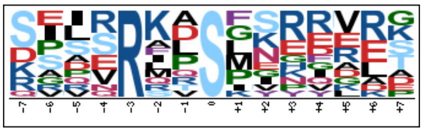

**C.**

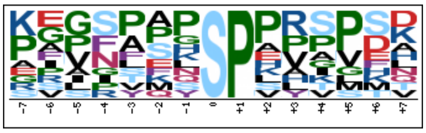

**Supplemental Figure S2.** Motif-X analysis of phosphopeptides with decreased abundance in *mlk1/2/3* mutants.

Phosphopeptides with altered abundance in *mlk1/2/3* (ZT12) mutants where extended and aligned using PEPTIDEXTENDER (http://schwartzlab.uconn.edu/pepextend). Motif-X analysis was then preformed with the probability threshold was set to p-value ≤ $10^{-6}$, the occurrence threshold was set to 10. The default IPI Arabidopsis Proteome data set was used as the background data set.

Supplemental Table S1. Summary of Pairwise Comparisons

| | Global Proteome | | | Phosphoproteome | | |
|---|---|---|---|---|---|---|
| Pairwise Comparison | Total Peptides with Altered Abundance | Proteins with Increase Abundance | Proteins with Decrease Abundance | Total Peptides with Altered Abundance | Proteins with Increase Abundance | Proteins with Decrease Abundance |
| *mlk123*_ZT12/ WT_ZT12 | 15 | 0 | 13 | 113 | 44 | 50 |
| *mlk134*_ZT12/ WT_ZT12 | 232 | 28 | 95 | 429 | 103 | 190 |
| *mlk123*_ZT14/ WT_ZT14 | 20 | 6 | 7 | 170 | 87 | 63 |
| *mlk134*_ZT14/ WT_ZT14 | 228 | 76 | 36 | 306 | 108 | 115 |

**Supplemental Table S2. REVIGO Cellular Component GO-term Enrichment Analysis**

***mlk1/2/3* ZT12**

| GO-term ID | Description | Frequency | $Log_{10}$ p-value | Uniqueness | Dispensability |
|---|---|---|---|---|---|
| **GO:0005654** | **nucleoplasm** | **1.37%** | **-3.0182** | **0.15** | **0** |
| GO:0044451 | *nucleoplasm part* | *0.90%* | *-2.0482* | *0.17* | *0.82* |
| **GO:0043226** | **organelle** | **20.79%** | **-2.5969** | **0.75** | **0** |
| **GO:0005634** | **nucleus** | **8.97%** | **-4.4283** | **0.13** | **0.51** |
| GO:0043231 | *intracellular membrane-bounded organelle* | *13.76%* | *-2.6364* | *0.12* | *0.75* |
| **GO:0043227** | **membrane-bounded organelle** | **14.45%** | **-3.382** | **0.17** | **0.57** |
| GO:0043229 | *intracellular organelle* | *19.92%* | *-2.6144* | *0.16* | *0.7* |

***mlk1/3/4* ZT12**

| GO-term ID | Description | Frequency | $Log_{10}$ p-value | Uniqueness | Dispensability |
|---|---|---|---|---|---|
| **GO:0005575** | **cellular_component** | **100.00%** | **-1.7986** | **1** | **0** |
| **GO:0005623** | **cell** | **53.55%** | **-12.1624** | **0.97** | **0** |
| **GO:0005654** | **nucleoplasm** | **1.37%** | **-13.8069** | **0.31** | **0** |
| GO:0044451 | *nucleoplasm part* | *0.90%* | *-10.1113* | *0.33* | *0.82* |
| GO:0070013 | *intracellular organelle lumen* | *2.74%* | *-8.9747* | *0.3* | *0.95* |
| GO:0031981 | *nuclear lumen* | *2.29%* | *-10.58* | *0.29* | *0.87* |
| GO:0043233 | *organelle lumen* | *2.74%* | *-8.9747* | *0.33* | *0.97* |
| **GO:0031974** | **membrane-enclosed lumen** | **2.74%** | **-8.9747** | **0.94** | **0** |
| **GO:0032991** | **macromolecular complex** | **14.01%** | **-3.6517** | **0.95** | **0** |
| **GO:0043226** | **organelle** | **20.79%** | **-11.9788** | **0.95** | **0** |
| **GO:0043234** | **protein complex** | **6.42%** | **-4.0467** | **0.88** | **0** |
| **GO:0005829** | **cytosol** | **2.55%** | **-6.6478** | **0.72** | **0.17** |
| **GO:0044424** | **intracellular part** | **35.65%** | **-11.7077** | **0.68** | **0.22** |
| **GO:0005622** | **intracellular** | **41.18%** | **-11.4711** | **0.77** | **0.32** |
| **GO:0044444** | **cytoplasmic part** | **12.66%** | **-2.011** | **0.66** | **0.35** |
| **GO:0044464** | **cell part** | **52.39%** | **-10.7959** | **0.77** | **0.38** |
| **GO:0043227** | **membrane-bounded organelle** | **14.45%** | **-12.9172** | **0.47** | **0.4** |

| | | | | | |
|---|---|---|---|---|---|
| GO:0043229 | *intracellular organelle* | *19.92%* | *-11.5935* | *0.39* | *0.7* |
| **GO:0000785** | **chromatin** | **0.47%** | **-9.0726** | **0.41** | **0.46** |
| GO:0044427 | *chromosomal part* | *1.12%* | *-7.4962* | *0.39* | *0.84* |
| GO:0000790 | *nuclear chromatin* | *0.23%* | *-1.6216* | *0.34* | *0.78* |
| **GO:0005694** | **chromosome** | **1.51%** | **-6.6556** | **0.5** | **0.49** |
| **GO:0005652** | **nuclear lamina** | **0.00%** | **-2.2358** | **0.53** | **0.51** |
| **GO:0043228** | **non-membrane-bounded organelle** | **8.41%** | **-2.4597** | **0.5** | **0.56** |
| **GO:0044428** | **nuclear part** | **3.12%** | **-10.7447** | **0.37** | **0.57** |
| GO:0044446 | *intracellular organelle part* | *8.94%* | *-7.9355* | *0.32* | *0.73* |
| **GO:0044422** | **organelle part** | **9.43%** | **-7.8861** | **0.5** | **0.58** |
| **GO:0005634** | **nucleus** | **8.97%** | **-19.4547** | **0.39** | **0.58** |
| GO:0043231 | *intracellular membrane-bounded organelle* | *13.76%* | *-12.6216* | *0.37* | *0.75* |
| **GO:0031965** | **nuclear membrane** | **0.10%** | **-1.7721** | **0.46** | **0.6** |
| **GO:0005849** | **mRNA cleavage factor complex** | **0.05%** | **-1.71** | **0.42** | **0.62** |
| **GO:0016607** | **nuclear speck** | **0.09%** | **-3.6498** | **0.42** | **0.65** |
| GO:0016604 | *nuclear body* | *0.19%* | *-2.4895* | *0.39* | *0.7* |
| **GO:0043232** | **intracellular non-membrane-bounded organelle** | **7.95%** | **-2.4597** | **0.41** | **0.67** |

***mlk1/2/3* ZT14**

| GO-term ID | Description | Frequency | Log$_{10}$ p-value | Uniqueness | Dispensability |
|---|---|---|---|---|---|
| **GO:0005623** | **cell** | **53.55%** | **-1.3298** | **0.93** | **0** |
| **GO:0031974** | **membrane-enclosed lumen** | **2.74%** | **-1.5258** | **0.87** | **0** |
| **GO:0044427** | **chromosomal part** | **1.12%** | **-3.9547** | **0.26** | **0** |
| GO:0000785 | *chromatin* | *0.47%* | *-1.9872* | *0.27* | *0.84* |
| GO:1902494 | catalytic complex | 3.73% | -3.6925 | 0.82 | 0 |
| **GO:0044422** | **organelle part** | **9.43%** | **-1.5686** | **0.35** | **0.36** |
| **GO:0043228** | **non-membrane-bounded organelle** | **8.41%** | **-1.3565** | **0.36** | **0.51** |
| **GO:0031981** | **nuclear lumen** | **2.29%** | **-2.757** | **0.23** | **0.53** |
| GO:0043233 | *organelle lumen* | *2.74%* | *-1.5258* | *0.24* | *0.97* |
| GO:0070013 | *intracellular organelle lumen* | *2.74%* | *-1.5258* | *0.23* | *0.95* |

| | | | | | |
|---|---|---|---|---|---|
| **GO:0005694** | **chromosome** | **1.51%** | **-2.9914** | **0.37** | **0.54** |
| **GO:0005634** | **nucleus** | **8.97%** | **-2.8239** | **0.32** | **0.55** |
| **GO:0044428** | **nuclear part** | **3.12%** | **-2.063** | **0.27** | **0.61** |
| GO:0044446 | *intracellular organelle part* | *8.94%* | *-1.5918* | *0.22* | *0.73* |
| **GO:0043232** | **intracellular non-membrane-bounded organelle** | **7.95%** | **-1.3565** | **0.28** | **0.67** |

***mlk1/3/4* ZT14**

| GO-term ID | Description | Frequency | $Log_{10}$ p-value | Uniqueness | Dispensability |
|---|---|---|---|---|---|
| **GO:0005623** | **cell** | **53.55%** | **-5.2218** | **0.95** | **0** |
| **GO:0005654** | **nucleoplasm** | **1.37%** | **-6.6778** | **0.29** | **0** |
| GO:0043233 | *organelle lumen* | *2.74%* | *-2.4559* | *0.31* | *0.97* |
| GO:0044451 | *nucleoplasm part* | *0.90%* | *-4.9706* | *0.31* | *0.82* |
| GO:0070013 | *intracellular organelle lumen* | *2.74%* | *-2.4559* | *0.27* | *0.95* |
| GO:0031981 | *nuclear lumen* | *2.29%* | *-2.9547* | *0.26* | *0.87* |
| **GO:0031974** | **membrane-enclosed lumen** | **2.74%** | **-2.4559** | **0.91** | **0** |
| **GO:0043226** | **organelle** | **20.79%** | **-4.0615** | **0.92** | **0** |
| **GO:0044464** | **cell part** | **52.39%** | **-4.4935** | **0.72** | **0.12** |
| **GO:0005829** | **cytosol** | **2.55%** | **-3.8268** | **0.67** | **0.17** |
| **GO:0044424** | **intracellular part** | **35.65%** | **-3.8386** | **0.62** | **0.35** |
| **GO:0005622** | **intracellular** | **41.18%** | **-3.52** | **0.71** | **0.38** |
| **GO:0044427** | **chromosomal part** | **1.12%** | **-3.1024** | **0.36** | **0.5** |
| GO:0000785 | *chromatin* | *0.47%* | *-1.8861* | *0.4* | *0.84* |
| **GO:0070449** | **elongin complex** | **0.01%** | **-1.719** | **0.48** | **0.53** |
| **GO:0005694** | **chromosome** | **1.51%** | **-2.1232** | **0.47** | **0.54** |
| **GO:0044428** | **nuclear part** | **3.12%** | **-1.857** | **0.33** | **0.57** |
| **GO:0005634** | **nucleus** | **8.97%** | **-8.9136** | **0.35** | **0.58** |
| GO:0043231 | *intracellular membrane-bounded organelle* | *13.76%* | *-3.3686* | *0.33* | *0.75* |
| **GO:0043229** | **intracellular organelle** | **19.92%** | **-4.091** | **0.35** | **0.62** |
| GO:0043227 | *membrane-bounded organelle* | *14.45%* | *-3.6861* | *0.44* | *0.7* |

**Supplemental Table S3. REVIGO Biological Process GO-term Enrichment Analysis**

| | | *mlk1/2/3* ZT12 | | | |
|---|---|---|---|---|---|
| **GO-term ID** | **Description** | **Frequency** | **$Log_{10}$ p-value** | **Uniqueness** | **Dispensability** |
| **GO:0040029** | **regulation of gene expression, epigenetic** | **0.13%** | **-1.4572** | **1** | **0** |

| | | *mlk1/3/4* ZT12 | | | |
|---|---|---|---|---|---|
| **GO-term ID** | **Description** | **Frequency** | **$Log_{10}$ p-value** | **Uniqueness** | **Dispensability** |
| **GO:0006325** | **chromatin organization** | **0.67%** | **-10.4078** | **0.88** | **0** |
| GO:0016043 | *cellular component organization* | *7.24%* | *-2.5575* | *0.86* | *0.75* |
| GO:0007064 | *mitotic sister chromatid cohesion* | *0.05%* | *-2.1267* | *0.89* | *0.54* |
| GO:0051276 | *chromosome organization* | *1.48%* | *-10.1433* | *0.87* | *0.53* |
| GO:0016570 | *histone modification* | *0.37%* | *-2.0696* | *0.77* | *0.9* |
| GO:0016569 | *covalent chromatin modification* | *0.42%* | *-4.2168* | *0.8* | *0.7* |
| GO:0006996 | *organelle organization* | *3.60%* | *-4.5768* | *0.86* | *0.65* |
| **GO:0007623** | **circadian rhythm** | **0.06%** | **-1.6968** | **0.98** | **0** |
| **GO:0009987** | **cellular process** | **63.78%** | **-1.6536** | **0.99** | **0** |
| **GO:0010033** | **response to organic substance** | **0.90%** | **-4.1475** | **0.76** | **0** |
| GO:0001101 | *response to acid chemical* | *0.12%* | *-1.5017* | *0.79* | *0.59* |
| GO:1901701 | *cellular response to oxygen-containing compound* | *0.35%* | *-2.6778* | *0.71* | *0.8* |
| GO:1901700 | *response to oxygen-containing compound* | *0.50%* | *-1.9101* | *0.77* | *0.68* |
| GO:0097305 | *response to alcohol* | *0.06%* | *-3.15* | *0.76* | *0.55* |
| GO:0009737 | *response to abscisic acid* | *0.03%* | *-3.224* | *0.77* | *0.67* |
| GO:0009725 | *response to hormone* | *0.34%* | *-2.3585* | *0.74* | *0.79* |
| GO:0071359 | *cellular response to dsRNA* | *0.02%* | *-1.8794* | *0.73* | *0.81* |
| GO:0033993 | *response to lipid* | *0.23%* | *-2.6289* | *0.74* | *0.77* |
| GO:0070887 | *cellular response to chemical stimulus* | *1.01%* | *-2.4437* | *0.72* | *0.73* |
| GO:0043331 | *response to dsRNA* | *0.03%* | *-1.8794* | *0.76* | *0.83* |
| GO:0071407 | *cellular response to organic cyclic compound* | *0.17%* | *-2.6615* | *0.7* | *0.67* |
| GO:0071310 | *cellular response to organic substance* | *0.65%* | *-1.9393* | *0.69* | *0.87* |

| | | | | | |
|---|---|---|---|---|---|
| GO:0014070 | *response to organic cyclic compound* | *0.23%* | *-2.821* | *0.74* | *0.75* |
| **GO:0010605** | **negative regulation of macromolecule metabolic process** | **1.17%** | **-5.5421** | **0.51** | **0** |
| GO:0031050 | *dsRNA fragmentation* | *0.02%* | *-1.9355* | *0.59* | *0.98* |
| GO:0031047 | *gene silencing by RNA* | *0.09%* | *-4.1379* | *0.56* | *0.75* |
| GO:0048523 | *negative regulation of cellular process* | *1.83%* | *-2.0635* | *0.57* | *0.92* |
| GO:0031324 | *negative regulation of cellular metabolic process* | *1.17%* | *-3.3862* | *0.52* | *0.97* |
| GO:0031327 | *negative regulation of cellular biosynthetic process* | *0.77%* | *-4.1463* | *0.51* | *0.99* |
| GO:0035194 | *posttranscriptional gene silencing by RNA* | *0.03%* | *-3.3893* | *0.59* | *0.84* |
| GO:0035196 | *production of miRNAs involved in gene silencing by miRNA* | *0.01%* | *-2.5287* | *0.41* | *0.94* |
| GO:0035195 | *gene silencing by miRNA* | *0.02%* | *-2.8097* | *0.56* | *0.97* |
| GO:0051253 | *negative regulation of RNA metabolic process* | *0.63%* | *-1.5129* | *0.47* | *0.96* |
| GO:0070918 | *production of small RNA involved in gene silencing by RNA* | *0.02%* | *-1.9355* | *0.41* | *0.98* |
| GO:2000113 | *negative regulation of cellular macromolecule biosynthetic process* | *0.72%* | *-3.4461* | *0.49* | *0.98* |
| GO:0045934 | *negative regulation of nucleobase-containing compound metabolic process* | *0.70%* | *-1.8539* | *0.49* | *0.92* |
| GO:0010629 | *negative regulation of gene expression* | *0.78%* | *-5.1073* | *0.51* | *0.93* |
| GO:0009892 | *negative regulation of metabolic process* | *1.26%* | *-4.8962* | *0.57* | *0.87* |
| GO:0009890 | *negative regulation of biosynthetic process* | *0.77%* | *-4.0535* | *0.54* | *0.93* |
| GO:0010558 | *negative regulation of macromolecule biosynthetic process* | *0.74%* | *-4.4935* | *0.5* | *0.92* |
| GO:0016458 | *gene silencing* | *0.17%* | *-4.8125* | *0.55* | *0.8* |
| GO:0016441 | *posttranscriptional gene silencing* | *0.03%* | *-2.7447* | *0.59* | *0.84* |
| GO:0051172 | *negative regulation of nitrogen compound metabolic process* | *0.79%* | *-3.8477* | *0.53* | *0.93* |
| **GO:0032502** | **developmental process** | **2.81%** | **-2.6946** | **0.98** | **0** |

| | | | | | |
|---|---|---|---|---|---|
| **GO:0048511** | **rhythmic process** | **0.08%** | **-1.4271** | **0.98** | **0** |
| **GO:0048856** | **anatomical structure development** | **2.54%** | **-2.4547** | **0.97** | **0** |
| GO:0048731 | *system development* | *1.26%* | *-1.342* | *0.95* | *0.86* |
| **GO:0050896** | **response to stimulus** | **12.21%** | **-4.4584** | **0.98** | **0** |
| **GO:0065007** | **biological regulation** | **20.50%** | **-8.3458** | **0.99** | **0** |
| **GO:0071840** | **cellular component organization or biogenesis** | **8.57%** | **-1.5986** | **0.98** | **0** |
| **GO:0046483** | **heterocycle metabolic process** | **29.66%** | **-5.9066** | **0.85** | **0.05** |
| **GO:0006807** | **nitrogen compound metabolic process** | **38.74%** | **-2.7423** | **0.94** | **0.06** |
| **GO:1901360** | **organic cyclic compound metabolic process** | **30.32%** | **-4.8928** | **0.89** | **0.08** |
| **GO:0006396** | **RNA processing** | **3.21%** | **-4.2725** | **0.68** | **0.14** |
| **GO:0018193** | **peptidyl-amino acid modification** | **1.50%** | **-2.2958** | **0.8** | **0.18** |
| **GO:0043170** | **macromolecule metabolic process** | **39.49%** | **-2.4622** | **0.88** | **0.21** |
| **GO:0010467** | **gene expression** | **19.67%** | **-5.3325** | **0.81** | **0.22** |
| **GO:0006725** | **cellular aromatic compound metabolic process** | **29.63%** | **-5.3325** | **0.85** | **0.24** |
| **GO:0097298** | **regulation of nucleus size** | **0.00%** | **-2.8327** | **0.82** | **0.28** |
| **GO:0048519** | **negative regulation of biological process** | **1.98%** | **-3.7352** | **0.74** | **0.32** |
| **GO:0048518** | **positive regulation of biological process** | **1.74%** | **-2.1864** | **0.74** | **0.33** |
| **GO:0040029** | **regulation of gene expression, epigenetic** | **0.13%** | **-4.4157** | **0.68** | **0.34** |
| **GO:0006397** | **mRNA processing** | **0.56%** | **-4.266** | **0.7** | **0.36** |
| GO:0008380 | *RNA splicing* | *0.41%* | *-2.5214* | *0.71* | *0.62* |
| GO:0000377 | *RNA splicing, via transesterification reactions with bulged adenosine as nucleophile* | *0.32%* | *-1.8729* | *0.71* | *0.99* |
| GO:0000375 | *RNA splicing, via transesterification reactions* | *0.32%* | *-1.7721* | *0.71* | *0.95* |
| GO:0000398 | *mRNA splicing, via spliceosome* | *0.32%* | *-2.266* | *0.7* | *0.88* |
| **GO:0009719** | **response to endogenous stimulus** | **0.53%** | **-2.1871** | **0.83** | **0.37** |
| **GO:0016071** | **mRNA metabolic process** | **0.80%** | **-3.0283** | **0.75** | **0.37** |

| | | | | | |
|---|---|---|---|---|---|
| **GO:0090304** | **nucleic acid metabolic process** | **21.45%** | **-5.9706** | **0.66** | **0.39** |
| GO:0032774 | *RNA biosynthetic process* | *10.93%* | *-3.317* | *0.61* | *0.6* |
| GO:0018130 | *heterocycle biosynthetic process* | *17.39%* | *-2.9957* | *0.71* | *0.52* |
| GO:0016070 | *RNA metabolic process* | *15.95%* | *-5.0969* | *0.65* | *0.58* |
| GO:0006351 | *transcription, DNA-templated* | *10.66%* | *-2.7986* | *0.58* | *0.74* |
| GO:0097659 | *nucleic acid-templated transcription* | *10.72%* | *-3.0814* | *0.6* | *0.88* |
| GO:0006139 | *nucleobase-containing compound metabolic process* | *26.55%* | *-5.9747* | *0.7* | *0.6* |
| GO:0019438 | *aromatic compound biosynthetic process* | *16.95%* | *-2.7258* | *0.72* | *0.52* |
| GO:1901362 | *organic cyclic compound biosynthetic process* | *17.87%* | *-2.5017* | *0.73* | *0.51* |
| GO:0034654 | *nucleobase-containing compound biosynthetic process* | *14.53%* | *-2.7167* | *0.66* | *0.71* |
| **GO:0048583** | **regulation of response to stimulus** | **1.12%** | **-2.6925** | **0.63** | **0.4** |
| **GO:0044260** | **cellular macromolecule metabolic process** | **34.28%** | **-3.4685** | **0.74** | **0.43** |
| **GO:0034641** | **cellular nitrogen compound metabolic process** | **34.14%** | **-4.1543** | **0.77** | **0.44** |
| **GO:0050789** | **regulation of biological process** | **19.37%** | **-7.8794** | **0.66** | **0.45** |
| GO:0080090 | *regulation of primary metabolic process* | *11.68%* | *-6.5952* | *0.55* | *0.87* |
| GO:0010468 | *regulation of gene expression* | *10.82%* | *-5.2541* | *0.49* | *0.86* |
| GO:0031323 | *regulation of cellular metabolic process* | *11.66%* | *-4.2907* | *0.52* | *0.87* |
| GO:0031326 | *regulation of cellular biosynthetic process* | *10.82%* | *-4.4377* | *0.49* | *0.89* |
| GO:0051252 | *regulation of RNA metabolic process* | *10.03%* | *-3.3936* | *0.42* | *0.86* |
| GO:2001141 | *regulation of RNA biosynthetic process* | *9.97%* | *-2.9469* | *0.41* | *0.9* |
| GO:0019219 | *regulation of nucleobase-containing compound metabolic process* | *10.26%* | *-3.7799* | *0.46* | *0.85* |
| GO:2000112 | *regulation of cellular macromolecule biosynthetic process* | *10.68%* | *-4.1681* | *0.45* | *0.85* |
| GO:0006355 | *regulation of transcription, DNA-templated* | *9.92%* | *-2.6576* | *0.41* | *0.9* |
| GO:0060255 | *regulation of macromolecule metabolic process* | *11.72%* | *-7.0004* | *0.51* | *0.67* |

| | | | | | |
|---|---|---|---|---|---|
| GO:0019222 | *regulation of metabolic process* | *11.94%* | *-5.8539* | *0.65* | *0.67* |
| GO:0050794 | *regulation of cellular process* | *18.84%* | *-5.0937* | *0.6* | *0.76* |
| GO:1903506 | *regulation of nucleic acid-templated transcription* | *9.97%* | *-2.9469* | *0.41* | *0.89* |
| GO:0009889 | *regulation of biosynthetic process* | *10.83%* | *-4.3382* | *0.53* | *0.86* |
| GO:0010556 | *regulation of macromolecule biosynthetic process* | *10.75%* | *-4.4401* | *0.48* | *0.89* |
| GO:0051171 | *regulation of nitrogen compound metabolic process* | *10.93%* | *-6.4908* | *0.52* | *0.86* |
| **GO:0042221** | **response to chemical** | **3.07%** | **-2.6162** | **0.81** | **0.46** |
| GO:0051716 | *cellular response to stimulus* | *9.56%* | *-3.384* | *0.74* | *0.63* |
| GO:0006281 | *DNA repair* | *2.23%* | *-1.4724* | *0.59* | *0.9* |
| GO:0033554 | *cellular response to stress* | *2.97%* | *-1.4737* | *0.75* | *0.81* |
| GO:0006974 | *cellular response to DNA damage stimulus* | *2.36%* | *-1.7122* | *0.75* | *0.51* |
| GO:0080134 | *regulation of response to stress* | *0.34%* | *-1.684* | *0.65* | *0.62* |
| **GO:0044271** | **cellular nitrogen compound biosynthetic process** | **22.50%** | **-1.7328** | **0.71** | **0.47** |
| **GO:0018205** | **peptidyl-lysine modification** | **0.36%** | **-1.3134** | **0.83** | **0.48** |

***mlk1/2/3* ZT14**

| GO-term ID | Description | Frequency | $Log_{10}$ p-value | Uniqueness | Dispensability |
|---|---|---|---|---|---|
| **GO:0051276** | **chromosome organization** | **1.48%** | **-2.7852** | **0** | **0** |
| GO:0007064 | *mitotic sister chromatid cohesion* | *0.05%* | *-1.7645* | *0* | *0.54* |
| GO:0006325 | *chromatin organization* | *0.67%* | *-1.3072* | *0* | *0.53* |

***mlk1/3/4* ZT14**

| GO-term ID | Description | Frequency | $Log_{10}$ p-value | Uniqueness | Dispensability |
|---|---|---|---|---|---|
| **GO:0009628** | **response to abiotic stimulus** | **0.57%** | **-1.857** | **0.91** | **0** |
| **GO:0009987** | **cellular process** | **63.78%** | **-2.1512** | **0.97** | **0** |
| **GO:0010629** | **negative regulation of gene expression** | **0.78%** | **-1.644** | **0.62** | **0** |
| **GO:0050896** | **response to stimulus** | **12.21%** | **-1.8962** | **0.94** | **0** |

| | | | | | |
|---|---|---|---|---|---|
| **GO:0051276** | **chromosome organization** | **1.48%** | **-2.1175** | **0.86** | **0** |
| GO:0006325 | *chromatin organization* | *0.67%* | *-1.308* | *0.86* | *0.53* |
| **GO:0065007** | **biological regulation** | **20.50%** | **-2.4425** | **0.94** | **0** |
| **GO:0009058** | **biosynthetic process** | **31.61%** | **-1.6655** | **0.88** | **0.02** |
| **GO:0044249** | **cellular biosynthetic process** | **30.05%** | **-2.762** | **0.65** | **0.06** |
| GO:1901576 | *organic substance biosynthetic process* | *30.37%* | *-2.2013* | *0.68* | *0.66* |
| GO:0018130 | *heterocycle biosynthetic process* | *17.39%* | *-1.7144* | *0.68* | *0.53* |
| GO:0019438 | *aromatic compound biosynthetic process* | *16.95%* | *-1.5114* | *0.68* | *0.53* |
| GO:1901362 | *organic cyclic compound biosynthetic process* | *17.87%* | *-1.752* | *0.7* | *0.54* |
| **GO:1901360** | **organic cyclic compound metabolic process** | **30.32%** | **-1.301** | **0.86** | **0.07** |
| **GO:0071704** | **organic substance metabolic process** | **58.36%** | **-1.5157** | **0.89** | **0.08** |
| **GO:0006807** | **nitrogen compound metabolic process** | **38.74%** | **-1.4377** | **0.88** | **0.09** |
| **GO:0044237** | **cellular metabolic process** | **53.06%** | **-2.2358** | **0.83** | **0.18** |
| **GO:0046483** | **heterocycle metabolic process** | **29.66%** | **-1.3439** | **0.81** | **0.25** |
| **GO:0097305** | **response to alcohol** | **0.06%** | **-1.5686** | **0.91** | **0.29** |
| **GO:0019222** | **regulation of metabolic process** | **11.94%** | **-3.5229** | **0.56** | **0.38** |
| GO:0050794 | *regulation of cellular process* | *18.84%* | *-1.5935* | *0.52* | *0.76* |
| GO:0050789 | *regulation of biological process* | *19.37%* | *-3.0804* | *0.57* | *0.67* |
| GO:0080090 | *regulation of primary metabolic process* | *11.68%* | *-2.6003* | *0.49* | *0.87* |
| GO:0009889 | *regulation of biosynthetic process* | *10.83%* | *-1.7986* | *0.4* | *0.86* |
| GO:0031323 | *regulation of cellular metabolic process* | *11.66%* | *-1.9957* | *0.45* | *0.87* |
| GO:0031326 | *regulation of cellular biosynthetic process* | *10.82%* | *-1.8665* | *0.38* | *0.86* |
| GO:0060255 | *regulation of macromolecule metabolic process* | *11.72%* | *-2.8069* | *0.47* | *0.62* |
| GO:0051171 | *regulation of nitrogen compound metabolic process* | *10.93%* | *-2.7959* | *0.49* | *0.86* |
